# A metabolic map of the DNA damage response identifies PRDX1 in nuclear ROS scavenging and aspartate synthesis

**DOI:** 10.1101/2022.08.01.500855

**Authors:** Amandine Moretton, Savvas Kourtis, Chiara Calabrò, Antoni Gañez Zapater, Frédéric Fontaine, André C. Müller, Joanna I. Loizou, Sara Sdelci

**Author notes:** Equal contribution (Amandine Moretton, Savvas Kourtis).

## Abstract

While cellular metabolism impacts the DNA damage response, a systematic understanding of the metabolic requirements that are crucial for DNA damage repair has yet to be reported. Here, we investigate the metabolic enzymes and processes that are essential when cells are exposed to DNA damage. By integrating functional genomics with chromatin proteomics and metabolomics, we provide a detailed description of the interplay between cellular metabolism and the DNA damage response. Subsequent analysis identified Peroxiredoxin 1, PRDX1, as fundamental for DNA damage repair. During the DNA damage response, PRDX1 translocates to the nucleus where it is required to reduce DNA damage-induced nuclear reactive oxygen species. Moreover, PRDX1 regulates aspartate availability, which is required for the DNA damage-induced upregulation of *de novo* nucleotide synthesis. Loss of PRDX1 leads to an impairment in the clearance of DNA damage, accumulation of replicative stress and cell proliferation defects, thus revealing a crucial role for PRDX1 as a DNA damage surveillance factor.

## Introduction

Maintaining genome integrity via repair of DNA damage is a key biological process required to suppress diseases, including growth retardation, malignancy, neurodegeneration and congenital anomalies (Jackson & Bartek, 2009). DNA is continually subjected to both exogenous and endogenous mutagens and hence cells have evolved distinct DNA repair mechanisms to counter different types of DNA damage (Hoeijmakers, 2001). In response to DNA damage, cells elicit a signaling cascade to repair the damaged DNA and/or arrest the cell cycle. The cascade results in activation of a specific repair machinery, which is recruited to the relevant site on chromatin. If the damage is beyond repair, sustained signaling from the damaged site might promote cells to enter senescence or undergo apoptosis.

Recent years have seen remarkable progress in unraveling the diverse mechanisms of the DNA damage response (DDR), broadening our knowledge of the diverse DDR pathways. Through such work, it has emerged that cellular metabolism not only generates DNA damage but also affects DNA repair (Moretton & Loizou, 2020; Turgeon *et al*, 2018). Metabolic reactions give rise to diverse types of DNA damage. Reactive oxygen species (ROS), mainly produced by oxidative phosphorylation, induce oxidative DNA damage, which is prevented by antioxidant metabolites such as glutathione (GSH) and nicotinamide adenine dinucleotide phosphate (NADPH) (Dizdaroglu, 1992; Harris *et al*, 2015). By-product metabolites such as aldehydes and alkylating agents can also form toxic adducts on DNA (Nakamura *et al*, 2014). Another aspect of the crosstalk between cellular metabolism and the DNA damage response is the tight control of the metabolic reactions involved in nucleotide synthesis. This is necessary for maintaining genomic integrity, both to avoid replication stress and nucleotide misincorporations, which result in DNA damage, and to ensure efficient DNA repair through the production of a local pool of nucleotides, in the vicinity of DNA double-strand breaks (DSBs) (Buckland *et al*, 2014; D’Angiolella *et al*, 2012). The function and recruitment of DNA repair enzymes to chromatin can additionally be regulated by metabolic enzymes and metabolites. For instance, the dealkylases AlkB homologs 2 and 3 (ALKBH2/3), which repair DNA adducts, use α-ketoglutaric acid (α-KG)—produced from glutamine—as a key substrate and are inhibited by the oncometabolite 2-hydroxyglutarate (2HG) (Tran *et al*, 2017; Wang *et al*, 2015). Finally, chromatin remodeling and epigenetic marks regulate the repair of DNA damage, especially DNA DSBs. Homologous recombination is promoted by histone acetylation, facilitated by the production of acetyl-CoA in the vicinity of DSBs (Sivanand *et al*, 2017). On the contrary, specific metabolites such as 2HG, fumarate, or succinate impair histone demethylation, preventing recruitment of homologous recombination factors by inhibiting the lysine specific demethylases 4A and 4B (KDM4A/B) (Sulkowski *et al*, 2020).

Yet, despite accumulating evidence of the dynamic interplay between metabolic factors and the DNA damage response, there has not been a systematic, unbiased study aimed at addressing how metabolic perturbations can affect DNA repair. Here we have identified the consequences of metabolic alterations on DNA damage and repair using a range of systematic approaches. Metabolism-focused CRISPR-Cas9 genetic screens, chromatin proteomics and targeted metabolomics following the induction of DNA damage using the chemotherapeutic topoisomerase 2 inhibitor etoposide revealed the aspects of metabolism that are crucial for the maintenance of genome integrity. Our findings indicated that nuclear ROS were generated following induction of DSBs by etoposide treatment and persisted 24 hours after release. Proteins of the Electron Transport Chain (ETC) were synthetic viable with etoposide and were recruited on chromatin. Additionally, the cellular metabolome was drastically perturbed by the DNA damaging treatment, especially nucleosides and nucleoside-related metabolites. The intersection of these datasets identified Peroxiredoxin 1, PRDX1, as a key enzyme of the DNA damage response, through regulation of ROS scavenging, control of aspartate metabolism and prevention of replication stress.

## Results

### Genetic map of metabolic factors that impact the DNA damage response

A thorough characterization of the DNA damage response-associated metabolic requirements has not yet been reported. Thus, to study the impact of metabolic alterations on the DNA damage response, we performed a CRISPR-Cas9 genetic screen to identify metabolic genes that affect cellular survival in response to DNA damage. Following transduction of the human cell line U2-OS with an sgRNA library targeting metabolic genes (Birsoy *et al*, 2015), we induced DSBs using etoposide, a common chemotherapeutic drug that inhibits topoisomerase II (Hande, 1998). Cells were exposed to etoposide for 3 hours at a concentration of 1μM followed by incubation in drug-free media for 24 hours, which allowed for clearance of DNA damage as shown by the restoration of γΗ2ΑΧ to basal levels (a DNA DSB marker (Sharma *et al*, 2012)) (Fig S1A). Cells were kept in culture for 5 days following etoposide treatment (denoted ‘survival CRISPR screen’) and untreated cells were cultured in parallel as a control (Fig 1A). DNA was extracted from both treated and untreated cells to identify depleted/enriched sgRNAs (Fig S1B-D, Source Data_1). As part of the data analysis, a cell cycle normalization step was performed to compensate for cell cycle defects that might occur due to the etoposide treatment (Fig S1E-F). Depleted sgRNAs allowed for the identification of metabolic genes that are required for cell survival upon etoposide treatment (synthetic lethality), while accumulated sgRNAs allowed for identification of genes that are synthetic viable with etoposide treatment.

**Figure 1:**
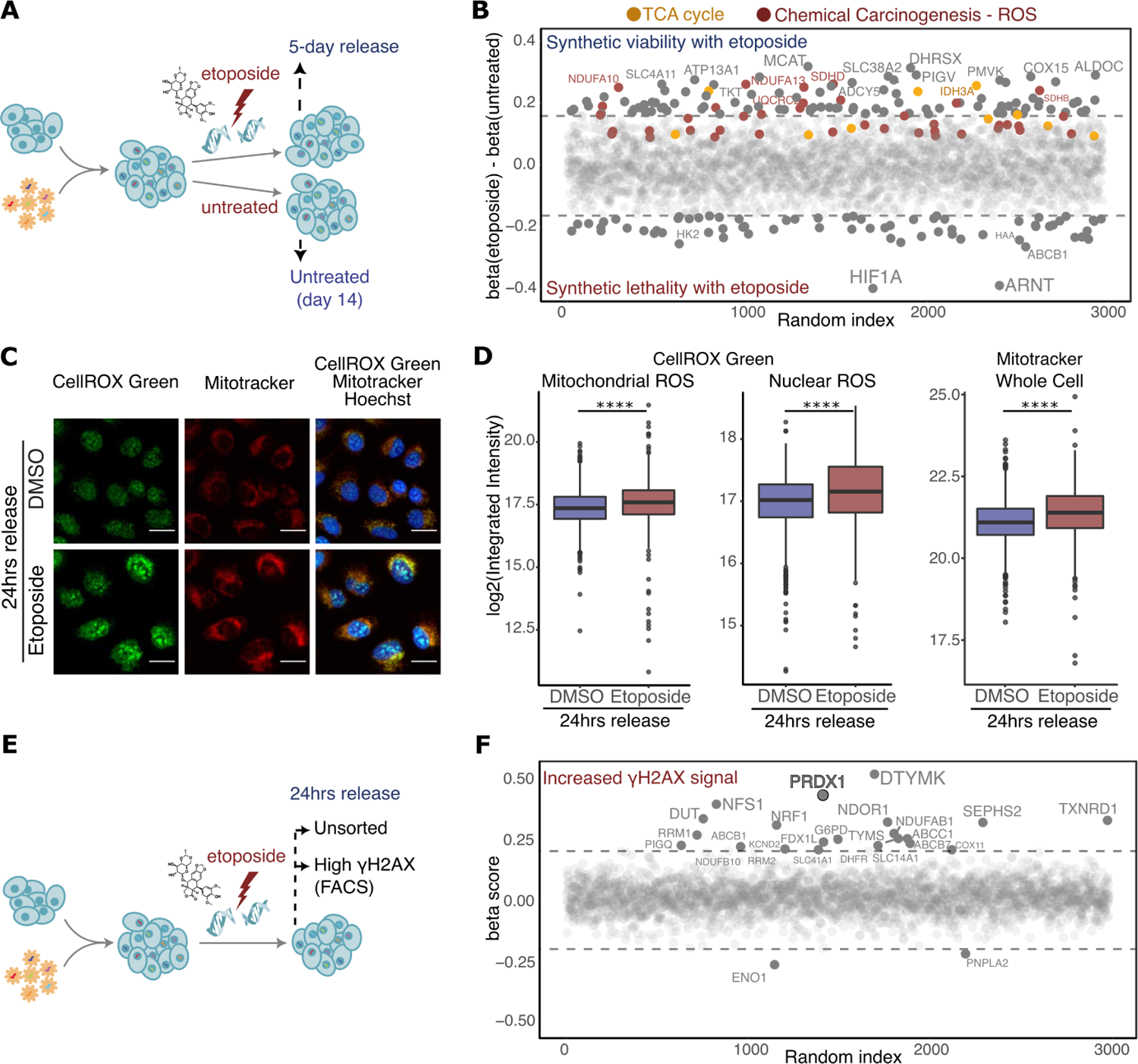
Metabolism-wide CRISPR-Cas9 screens identify ROS-related genes as synthetically viable with etoposide treatment. **(A)** Schematic representation of the etoposide survival CRISPR-Cas9 screen. **(B)** Genes synthetic lethal with etoposide survival are represented by negative β scores. Genes contributing to significant enrichment of KEGG terms are coloured. Sizes of the labels represent relative significance of screen hits. **(C)** Visualization of ROS (CellROX Green, in green) and mitochondria (Mitotracker, in red) within Hoechst-stained nuclei (in blue) in U2-OS WT cells in DMSO treated and 24 hours etoposide release conditions. Images were acquired on an Operetta High Content Screening System in confocal mode, scale bar is 25μm. **(D)** Quantification of images shown in (C), represented as log2 integrated intensity. A minimum of 1000 cells were quantified for each condition, using Harmony. P-values were calculated using linear regression on the log2 normalized values. **(E)** Schematic representation of the etoposide high-γΗ2ΑΧ CRISPR-Cas9 screen. **(F)** Genes necessary for γΗ2ΑΧ clearance are represented by positive β scores. Sizes of the labels represent relative significance of screen hits.

Hypoxia Inducible Factor 1 Subunit Alpha (HIF1A) and Aryl Hydrocarbon Receptor Nuclear Translocator (ARNT, also known as HIF1B), which interact to form the HIF1 heterodimeric transcription factor that promotes pro-glycolytic transcriptional states (Kim *et al*, 2006), were identified as the strongest synthetic lethal candidates (Fig 1B). The formation of the HIF1A-HIF1B heterodimer depends on HIF1A stabilization, which is commonly driven by hypoxia (Semenza, 2007), accumulation of ROS (Movafagh *et al*, 2015), and nutrient deprivation (Nishimoto *et al*, 2014), among other conditions. It is noteworthy that a previous study demonstrated that HIF1A mediates etoposide resistance in hypoxia conditions (Hussein *et al*, 2006).

Conversely, unbiased KEGG-based gene set enrichment analysis (GSEA) revealed that many genes of the Tricarboxylic Acid Cycle (KEGG term *Citrate cycle (TCA cycle)*) and the ETC (KEGG term *Chemical Carcinogenesis - ROS*), which are essential for oxidative phosphorylation and cellular respiration, were synthetically viable with etoposide treatment (Kanehisa & Goto, 2000; Wu *et al*, 2021). It has been previously shown that etoposide treatment generates ROS, which contribute to the cytotoxicity of this drug and arise from increased mitochondrial mass and respiration (Shin *et al*, 2016). ROS are important signaling molecules (Sies & Jones, 2020) that are physiologically produced during oxygen-consuming reactions in the mitochondria due to leaking electrons in the ETC, which cause partial oxygen reduction into superoxide radicals that are converted into H_2_O_2_ and hydroxyl radicals (Giorgio *et al*, 2007). Increased mitochondrial mass and respiration can lead to increased ROS levels and HIF1A stabilization, which in turn leads to downregulation of mitochondrial respiration (Yao *et al*, 2019). Using a fluorogenic probe to measure DNA-associated ROS, we observed that even the low etoposide concentration used for the genetic screen increased mitochondrial ROS, especially 24 hours after etoposide release (Fig 1C-D and S1G). This effect was accompanied by mitochondrial mass increase, detected with Mitotracker, which was clearly significant at 24 hours post-etoposide release (Fig 1C-D) but already appreciable after etoposide treatment (Fig S1G). While the augmented mitochondrial ROS levels can be the direct consequence of the mitochondrial mass increase (Fig S1H), we also observed a significant increase in nuclear ROS, which was present after etoposide treatment and more pronounced at 24 hours post-etoposide release (Fig 1C-D and Fig S1G). Thus, the etoposide-induced increase in ROS levels provided a connection between the synthetic lethal and synthetic viable genes in the screen. Taken together, the results of this genetic screen indicate that cells with a heightened glycolytic phenotype can better tolerate DNA damage, and this information may be important when considering anticancer regimens.

We reasoned that 5 days of release post-etoposide treatment would hamper the identification of metabolic genes that function early in the DDR, especially because the increase of cellular ROS was clearly observed at 24 hours post-etoposide release. Thus, we established a functional readout for identifying genes that affected levels of DNA damage by staining etoposide-treated cells for γΗ2ΑΧ levels and sorting the brightest population after 24 hours of etoposide release (denoted ‘high-γΗ2ΑΧ CRISPR screen’; Fig 1E and S1I). Sorting this population would identify genes that function in the resolution of DNA damage and thus the clearance of γΗ2ΑΧ. The quality control for this approach was performed as for the survival-CRISPR screen (Fig S1J-K).

Enolase 1 (ENO1) and Patatin Like Phospholipase Domain Containing 2 (PNPLA2), were the only two genes for which we found significantly depleted sgRNAs in the γΗ2ΑΧ bright population (Fig 1F, Source Data_1). This limited number of significant hits suggested that the lack of γΗ2ΑΧ clearance 24 hours post-DSBs induction did not depend on the enzymatic activity of any particular metabolic process. Supporting this hypothesis, it has been shown that ENO1 downregulation attenuates DNA damage induced by doxorubicin - a drug that intercalates within DNA, inhibiting the progression of topoisomerase II and leading to DSBs - independently of its enzymatic activity (Gao *et al*, 2015).

Conversely, when investigating sgRNA enrichment, we identified several genes belonging to the same metabolic pathway or sharing metabolic activity (Fig 1F). For example, multiple genes involved in nucleotide metabolism, such as Deoxythymidylate Kinase (DTYMK), Deoxyuridine Triphosphatase (DUT), Ribonucleotide Reductase Catalytic Subunit M1/2 (RRM1/2), Dihydrofolate Reductase (DHFR), and Thymidylate Synthetase (TYMS) belonged to this group. Depletion of these genes likely led to intracellular nucleotide imbalance and/or nucleotide shortage, which would result in nucleotide misincorporation, replication stress, and accumulation of DNA damage (Bester *et al*, 2011; Buckland *et al*., 2014). Moreover, sgRNAs targeting membrane transporters with known multidrug-resistance functions, like ATP Binding Cassette Subfamily C Member 1 (ABCC1), ATP Binding Cassette Subfamily C Member 7 (ABCB7), and ATP Binding Cassette Subfamily B Member 1 (ABCB1) also induced retention of γΗ2ΑΧ signal. Moreover, we observed that the depletion of genes with oxidoreductase activity, such as Peroxiredoxin 1 (PRDX1), Thioredoxin Reductase 1 (TXNRD1), NADPH Dependent Diflavin Oxidoreductase 1 (NDOR1), and Glucose-6-Phosphate Dehydrogenase (G6PD) which have a fundamental role in ROS balancing, was also associated with a lack of γΗ2ΑΧ clearance. In this group, PRDX1 displayed the most pronounced phenotype, indicating a strong connection between this enzyme and γΗ2ΑΧ clearance.

To validate the results of the high-γΗ2ΑΧ CRISPR screen, we performed an arrayed CRISPR screen using a library targeting the top genes whose depletion led to retention of γΗ2ΑΧ 24 hours after etoposide release. The radiomimetic compound neocarzinostatin (NCS) was used in parallel to etoposide. The concentration (60ng/mL) and time (1 hour) of NCS treatment allowed DNA damage clearance, as shown by the clearance of γΗ2ΑΧ staining following 20 hours of release (Fig S1L). γΗ2ΑΧ foci were quantified by microscopy in untreated, etoposide-, and NCS-treated cells immediately post-treatment or after 20 hours of release. As expected, targeting nucleotide metabolism-related genes strongly promoted the accumulation of γΗ2ΑΧ foci even in absence of exogenous DNA damage (Fig S1M). Targeting selected oxidoreductases (NDOR1, G6PD, TXNRD1, and PRDX1) did not induce a dramatic increase in baseline γΗ2ΑΧ foci but impeded the clearance of DNA damage 20 hours post-DSBs induction, indicating that these proteins might function in the DDR.

### Metabolic enzymes involved in DNA damage response localize on chromatin

24 hours post-etoposide release, we observed a marked increase in ROS within the cell nucleus (Fig 1C-D) Therefore, we addressed whether metabolic enzymes are associated with chromatin in response to DNA damage. To identify metabolic factors that may have a direct role in the DDR, we analyzed the chromatin-bound proteome of cells treated with etoposide. U2-OS were treated with DMSO or 1μM etoposide for 3 hours. Etoposide treated cells were released into drug-free media to allow for the monitoring of proteins bound to chromatin 24 hours post release (Fig 2A). Chromatome fractions were analyzed by mass-spectrometry (MS) and data analysis included batch correction (Fig S2A) and normalization (Fig S2B). The purity of the chromatomes was assessed by checking relative enrichment of protein in different cellular compartments, showing strong enrichment for chromatin-related proteins and a depletion in cytoplasmic and secretory proteins (Fig S2C-E).

**Figure 2:**
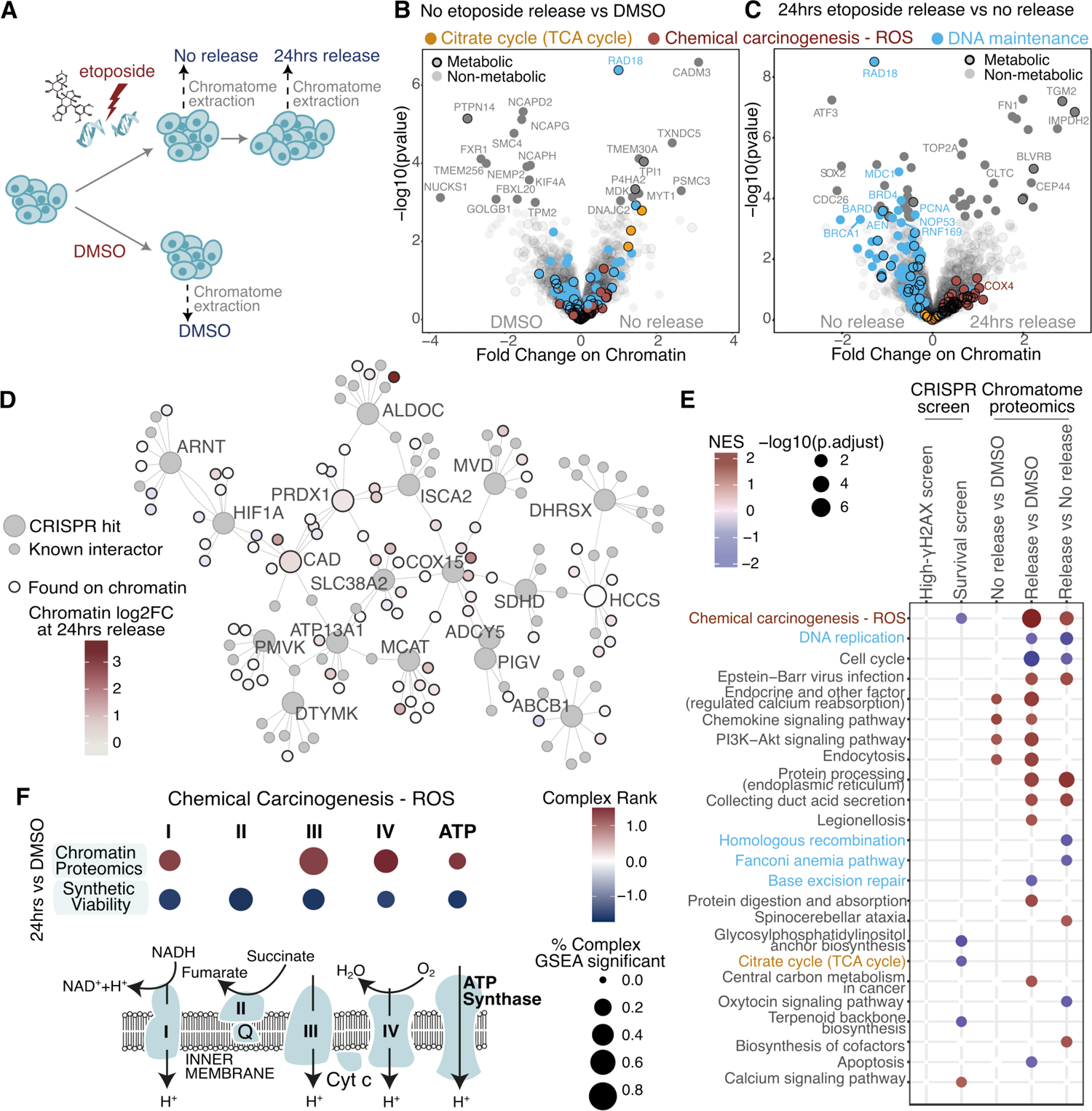
Chromatin proteomics reveals widespread accumulation of ROS-related genes on chromatin during etoposide release. **(A)** Schematic representation of etoposide treatment and release of U2-OS cells followed by chromatin extraction and DIA-MS acquisition. **(B, C)** Significant changes in protein abundance on chromatin upon etoposide treatment (B) compared to DMSO, and upon 24 hours release (C) compared to no release. Genes contributing to significant enrichment of KEGG terms are coloured. Genes are considered to have a metabolic function if they are either in the CRISPR metabolic library or in the Metabolic Atlas. **(D)** Protein-protein interaction network for top 5% gene hits in the CRISPR-Cas9 screens and their fold change on chromatin upon etoposide release. **(E)** Overlap of significant KEGG terms between the CRISPR-Cas9 screens and chromatin proteomics. **(F)** Mitochondrial electron transport chain genes significantly contributing to ‘Chemical Carcinogenesis - ROS’ KEGG term in chromatin proteomics and survival CRISPR screen.

In total, we identified 2,950 chromatin-bound proteins, of which 600 were metabolic factors, as annotated by the metabolic CRISPR library (Birsoy *et al*., 2015) and the Metabolic Atlas (Robinson *et al*, 2020) (Fig S2F, Table S1, Source Data_2). Eighty proteins were differently enriched or depleted on chromatin immediately after etoposide treatment (Fig 2B) and after 24 hours of etoposide release (Fig 2C). The chromatome composition remained altered 24 hours post release (Fig S2A), despite the strong reduction in the γΗ2ΑΧ signal at this time point (Fig S1A). This observation indicated that regardless of the γΗ2ΑΧ signal, 24 hours after etoposide release chromatin-associated alterations did not recover to their basal state. Several known DNA repair factors were differentially recruited to chromatin following etoposide treatment and release, thus validating our approach (Fig 2B-C and S2G). Amongst these, RAD18, which signals DNA damage and functions as an adaptor to recruit homologous recombination proteins (Huang *et al*, 2009), was enriched on chromatin following etoposide treatment and returned to basal levels after 24 hours release. BRCA1 and BARD1, which form a heterodimer involved in the DNA damage response to DSBs, followed a comparable recruitment pattern as RAD18 (Dai *et al*, 2021). DnaJ homolog subfamily C member 2 (DNAJC2) functions as a chromatin regulator. It binds monoubiquitylated histone H2A (Gracheva *et al*, 2016), an epigenetic mark that functions in DNA damage signaling and recruitment of DNA repair proteins early in the DDR, which explains its accumulation on chromatin immediately after etoposide treatment. On the contrary, Proliferating Cell Nuclear Antigen (PCNA), involved in DNA synthesis, and the cell cycle regulator Cell Division Cycle 26 (CDC26), which is required to elicit anaphase (Jin *et al*, 2008), were depleted from chromatin at 24 hours post-etoposide release, potentially due to a reduction of cellular proliferation and partial cell cycle arrest following DSB induction (Fig S2H). Nuclear casein kinase and cyclin-dependent kinase substrate 1 (NUCKS1), involved in homologous recombination, is lost from chromatin upon etoposide treatment and at 24 hours release. This transcription factor binds chromatin in a cell cycle-dependent manner and its levels increase in late G1, thus explaining this enrichment on chromatin (Hume *et al*, 2021; Parplys *et al*, 2015). Similarly, structural maintenance of chromosome 4 (SMC4), a component of the condensin complex facilitating the sister-chromatid condensation and mitosis, which is also involved in DNA repair (Wang & Wu, 2021), is depleted from chromatin immediately after etoposide treatment, but unlike NUCKS1, it is fully restored to basal levels 24 hours after release. Finally, Topoisomerase II alpha (TOP2A), the target of etoposide that forms covalent TOP2–DNA cleavage complexes, accumulated on chromatin upon etoposide release. In sum, chromatin-associated alteration of well-studied DDR proteins validated the robustness of our chromatome-DDR proteomics dataset.

Among the significantly altered proteins, 11 were metabolic enzymes (Table S1). Of note, 24 hours post-etoposide release, one of the chromatin-enriched metabolic enzymes was ENO1, whose sgRNAs were also found as depleted in the γΗ2ΑΧ-bright population, suggesting that the recruitment of this protein to chromatin may be important for DNA damage signaling. Additionally, we observed that several metabolic factors identified as differentially enriched or depleted in our genetic screens were found on chromatin (Table S2), or are known to have chromatin interactors (Fig 2D), suggesting that in response to DNA damage these proteins may have nuclear functions. Among them, we retrieved PRDX1, whose depletion resulted in the lack of γΗ2ΑΧ clearance 24 hours post-etoposide and 20 hours post-etoposide and NCS treatments (Fig 1F and S1L). This peroxiredoxin is a thiol-specific peroxidase that prevents the accumulation of ROS in cells and the generation of oxidative damage, thus functioning in H_2_O_2_ mediated signaling and cell growth upon oxidative stress (Neumann *et al*, 2009). Moreover, PRDX1 can also function as a molecular chaperone, when it forms a dimer after oxidation. The oligomer is then able to bind p53, c-Myc and other transcription factors, and regulate apoptosis and differentiation (Morinaka *et al*, 2011; Mu *et al*, 2002). We observed that the chromatin abundance of PRDX1 slightly increased following etoposide treatment, although not statistically significant (Fig 2D and S2I).

We performed KEGG-based GSEA with the datasets of the metabolism-focused CRISPR-Cas9 screens and chromatome proteomics (Fig 2E). The chromatome datasets indicated that DNA maintenance was still altered 24 hours after etoposide release, as highlighted by the enrichment of terms related to DNA replication, homologous recombination, Fanconi Anemia and base excision repair (grouped together under the term of *DNA maintenance* in Fig 2B-C). The *Chemical Carcinogenesis – ROS* term was shared between the survival CRISPR screen and the chromatome. The majority of genes responsible for the *Chemical Carcinogenesis – ROS* term were enzymes of the different ETC complexes, while no enzyme from complex 2 was detected on chromatin (Fig 2F and Table S3). We hypothesized that the chromatin localization of ETC enzymes may be related to the nuclear accumulation of ROS observed upon etoposide treatment (Fig 1C-D), which would explain why in the survival CRISPR screen the depletion of ETC proteins increased cell survival following etoposide withdrawal (Fig 1B). The accumulation of ETC complexes on chromatin was highest 24 hours after etoposide release (Fig 2C, S2J), which coincided with the high levels of nuclear ROS at the same time point (Fig 1D). This agrees with PRDX1 being a hit in the 24 hours ‘high-γΗ2ΑΧ CRISPR screen’ (Fig 1F), further supporting PRDX1 being involved in nuclear ROS scavenging.

We therefore further investigated the relationship between PRDX1, nuclear ROS and DNA damage. We reasoned that the chromatin localization of metabolic enzymes able to scavenge ROS, such as PRDX1, may be crucial to reduce intranuclear ROS levels. In line with this hypothesis, it has been reported that PRDX1-deficient cells specifically accumulate ROS in the nucleus (Egler *et al*, 2005). We observed that U2-OS PRDX1-depleted cells (U2-OS shPRDX1 cell population) showed a significant increase in nuclear ROS, a phenotype that was enhanced upon etoposide treatment and release (24 hours) (Fig 3A-B and S3A-C), suggesting the presence of a functional connection between the nuclear localization of PRDX1 and nuclear ROS accumulation. Additionally, we quantified γΗ2ΑΧ nuclear intensity together with PRDX1 nuclear intensity following etoposide treatment. We observed that γΗ2ΑΧ intensity increased immediately after treatment and returned to almost baseline levels at 20 hours of release (Fig 3C-D). In comparison, PRDX1 continued to accumulate in the nucleus (Fig 3C-D), probably due to its role in scavenging etoposide-induced nuclear ROS (Fig 1C-D). The nuclear accumulation of PRDX1 was associated with a small global increase in the expression of the enzyme (Fig S3A and D). Moreover, at 20 hours of etoposide release, there was a correlation between nuclear accumulation of PRDX1 and high levels of γΗ2ΑΧ (Pearson coefficient of 0.84, Fig S3E). Overall, these results indicated that PRDX1 nuclear accumulation correlates with γΗ2ΑΧ high levels, and as such may be required to scavenge etoposide-induced nuclear ROS.

**Figure 3:**
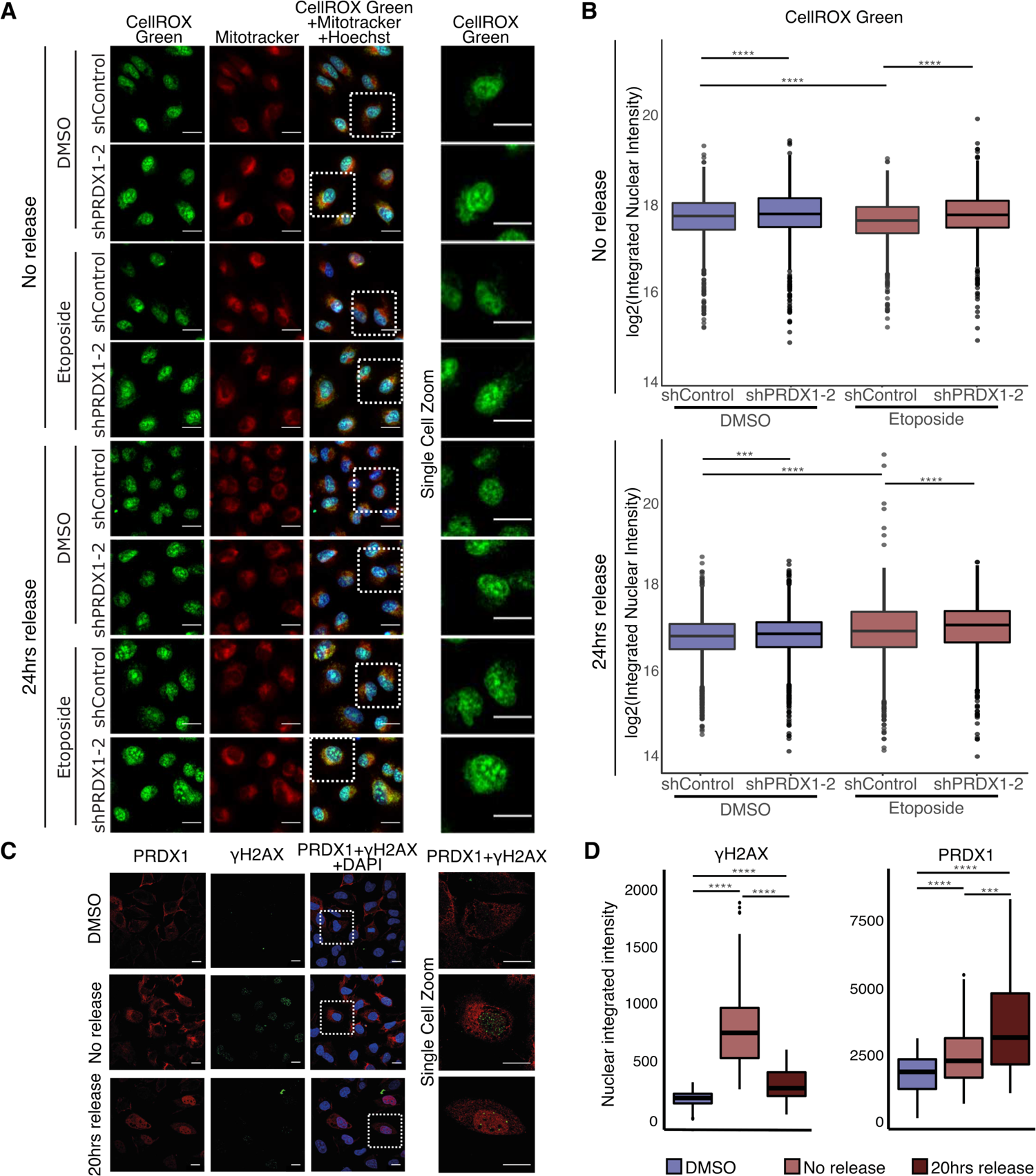
Nuclear ROS accumulates in the absence of PRDX1. **(A)** Visualization of ROS (CellROX Green, in green) and mitochondria (Mitotracker, in red) within Hoechst-stained nuclei (in blue) in U2-OS shControl and shPRDX1 cells at the indicated treatment conditions. Images were acquired with the Operetta High Content Screening System in confocal mode, scale bar is 25μm. **(B)** Quantification of images shown in (A), represented as log2 nuclear integrated intensity of CellROX Green immediately after etoposide treatment and at 24 hours release compared to DMSO control. A minimum of 1000 cells were quantified for each condition, using Harmony. P-values were calculated using linear regression on the log2 normalized values. **(C)** Visualization of PRDX1 (in red) and γΗ2ΑΧ (in green) within DAPI stained nuclei (in blue) in U2-OS WT cells at the indicated treatment conditions. Images were acquired on a confocal Zeiss LSM800 microscope, scale bar is 20μm. **(D)** Quantification of images shown in (C), represented as nuclear integrated intensities of γΗ2ΑΧ and PRDX1 signals. A minimum of 47 cells were quantified for each condition, using CellProfiler. P-values were calculated using the non-parametric Wilcoxon test.

### Metabolomics in the presence of DNA damage reveals PRDX1 to control aspartate synthesis

Since there is accumulating evidence that links metabolism and the DNA damage response, we assessed how the metabolic profile of cells is altered during the DDR. Therefore, we performed targeted metabolomics in U2-OS cells following etoposide treatment and release at different timepoints (Fig 4A). In total, 198 metabolites were measured, with a particular focus on nucleotide metabolism, amino acids and organic acids (Source Data_3). A total of 128 metabolites were detected in at least one sample, while 90 were consistently quantified in all samples (Fig S4A), with missing values following an intensity-dependent pattern (Fig S4B). The PCA plot showed a good clustering of biological replicates and indicated that, despite the reduction in the γΗ2ΑΧ signal (Fig S1A), the cellular metabolome remained altered at 24 hours etoposide release compared to the basal state (Fig S4C), in line with the findings from the chromatome dataset (Fig S2A).

**Figure 4:**
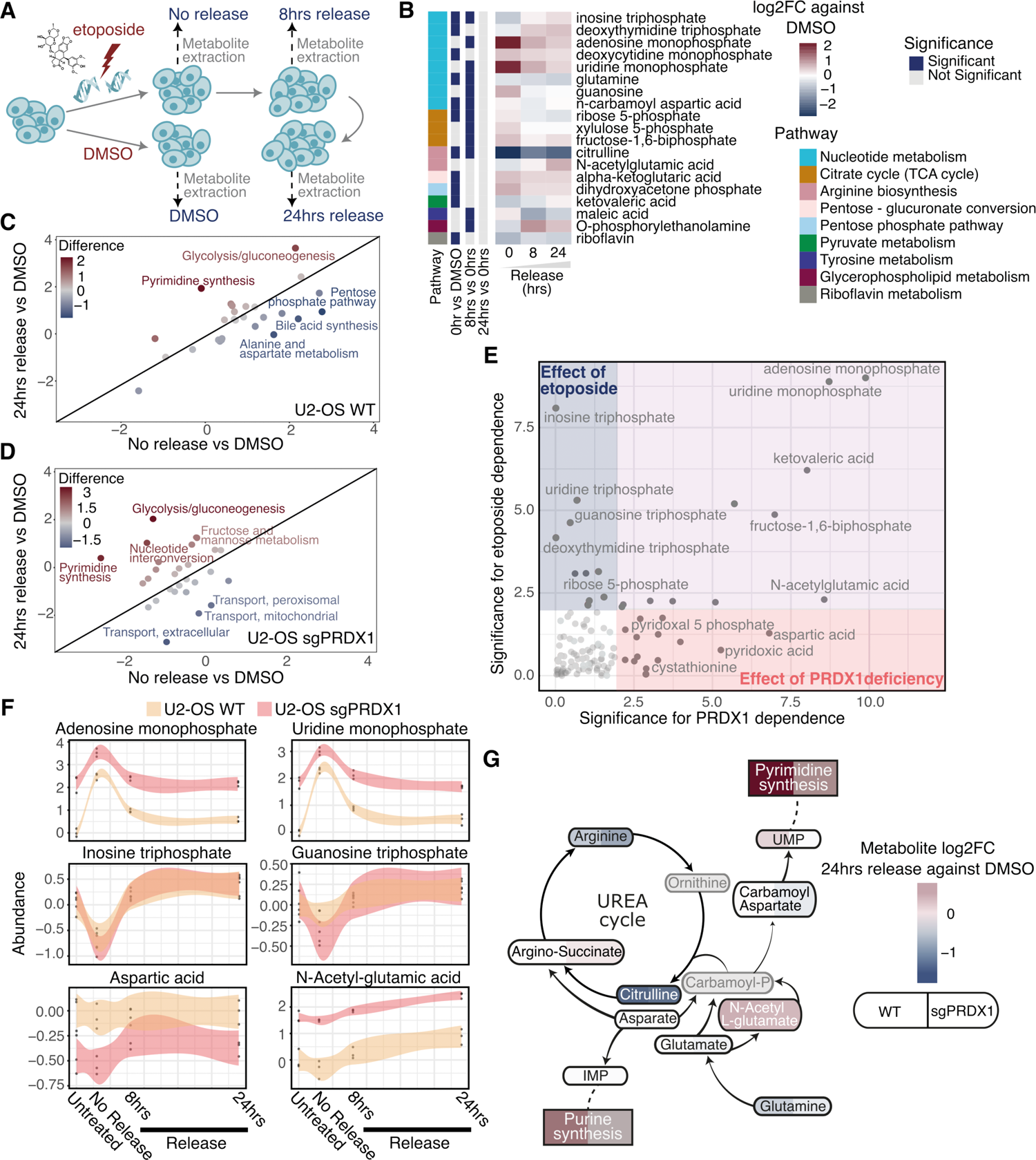
The cellular metabolome is drastically perturbed following etoposide treatment or PRDX1 loss. **(A)** Schematic representation of etoposide treatment and release of U2-OS cells followed by metabolite extraction and LC-MS/MS acquisition. **(B)** Relative abundances of metabolites that are significantly perturbed in at least one timepoint, represented as the log2 fold change compared to DMSO control. **(C)** Metabolic pathways altered in U2-OS WT cells at 24 hours etoposide release vs DMSO compared to no release vs DMSO. **(D)** Metabolic pathways altered in U2-OS sgPRDX1 cells at 24 hours etoposide release vs DMSO compared to no release vs DMSO. **(E)** PRDX1 deficiency-dependency and etoposide treatment-dependency of analyzed metabolites. **(F)** Abundance variations of example metabolites from (E) during the etoposide-relapse timecourse. **(G)** Relative abundances of key metabolites of the aspartate metabolism and connected metabolic pathways at 24 hours etoposide release compared to DMSO control, in U2-OS WT and sgPRDX1 cells.

When comparing DMSO and etoposide treated cells (Fig 4B), we observed that nucleosides and nucleoside-related metabolites were drastically perturbed. In particular, the identified triphosphate nucleosides decreased immediately after etoposide treatment, and increased at 8 and 24 hours release. This data suggested that nucleotides were acutely used upon DSB induction, probably as an outcome of repairing DNA damage, while during release into drug-free media the pools of nucleotides were replenished via *de novo* nucleotide synthesis. In agreement with this, monophosphate nucleoside levels were significantly increased at all time points but more pronounced immediately after etoposide treatment, suggesting that DSB induction and the activation of DDR triggered *de novo* nucleotide synthesis. Similarly, ribose and xylulose 5-phosphate, which are required for the synthesis of nucleoside sugar rings, rapidly increased following etoposide treatment and returned to basal levels during release into drug-free media. Successful *de novo* nucleotide synthesis additionally requires glutamine and aspartate. Specifically, aspartate alone is required for *de novo* purine synthesis (Pareek *et al*, 2021), while glutamine is the precursor of carbamoyl-phosphate, which, together with aspartate, is required for carbamoyl-aspartate production and *de novo* pyrimidine synthesis (Del Cano-Ochoa *et al*, 2019). Moreover, carbamoyl-phosphate and aspartate also contribute to citrulline and arginine synthesis (Shi *et al*, 2018). We did not record drastic changes in aspartate or carbamoyl-aspartate levels in our dataset, and carbamoyl-phosphate was not amongst the measured metabolites in our targeted approach. However, the reduction of citrulline and arginine levels observed at all given time points suggested that as an outcome of the DDR, aspartate and carbamoyl-phosphate are preferentially used for nucleotide synthesis.

The analysis of metabolites at the pathway level corroborated our observations. Nucleotide metabolism was affected during etoposide treatment, and remained upregulated after 24 hours of release, indicating once again that, despite the lack of γΗ2ΑΧ signal at this time point, cells were still affected by DNA damage. Conversely, the pentose phosphate pathway, which is required to synthesize the sugar backbone of nucleosides, was rapidly upregulated following etoposide treatment and returned to baseline levels after release (Fig 4C and S4D). Overall, the targeted metabolomics indicates that etoposide treatment activates *de novo* nucleotide synthesis and that 24 hours after etoposide release into drug-free media, the conversion from monophosphates to triphosphates is still active.

Given the fact that in our study PRDX1 was identified as having a central role in the DNA damage response, we addressed how loss of PRDX1 impacted the cellular metabolic state following DNA damage. Therefore, we performed targeted metabolomics comparing U2-OS wildtype (WT) with PRDX1-deficient (sgPRDX1) cell populations. In the principal component (PC) analysis, sgPRDX1 cells treated with DMSO overlapped with WT cells treated with etoposide, thus suggesting that loss of PRDX1 has an impact on the targeted metabolites, resembling metabolic changes observed following etoposide treatment (Fig S4C).

Our findings suggest that loss of PRDX1 broadly influences the metabolic state of cells even in the absence of exogenous DNA damage. When investigating significantly altered metabolites by comparing DMSO-treated and etoposide-treated sgPRDX1 cells, we detected overall minimal oscillation (Fig S4F), especially when compared with those observed in WT cells (Fig 4B). At the pathway level, we observed that during release in drug-free media, sgPRDX1 cells did not upregulate pyrimidine synthesis, as compared to WT cells (Fig 4 C-D). We next compared metabolic alterations due to etoposide treatment and/or PRDX1 deficiency and observed that monophosphate nucleotides were affected by both conditions while triphosphate nucleotides oscillations were strongly dependent on etoposide treatment. Aspartic acid and acetyl-glutamic acid showed a stronger dependency for PRDX1-loss rather than the etoposide treatment, suggesting the presence of a functional metabolic link between PRDX1 and aspartate metabolism (Fig 4E-F).

When investigating the dynamics of selected metabolite abundances, we observed that PRDX1-deficient cells had considerably lower levels of aspartate and acetyl-glutamic acid compared to WT cells, and that these marginally decreased following etoposide treatment. Overall, when comparing the metabolite changes observed at 24 hours release in the U2-OS WT and sgPRDX1 cells we observed that the latter displayed a stronger reduction of aspartate-dependent metabolites (arginine, citrulline and carbamoyl-aspartate), and failed to upregulate pyrimidine synthesis at the same level as WT cells (Fig 4G). This data suggested that the presence of PRDX1 is required to maintain baseline intracellular aspartate metabolism and that PRDX1 loss impairs the DDR in part due to a reduction in aspartate levels, altering aspartate-dependent metabolite production.

### Supplementation of aspartate partly rescues PRDX1 deficiency

Thus far, we hypothesize that PRDX1 has a bivalent role in the DDR by regulating nuclear ROS scavenging, and aspartate availability. Aspartate is a limiting metabolite for cancer cell proliferation in hypoxia and needs to be imported by active transport to sustain cellular growth upon inhibition of the mitochondrial electron transport chain (Garcia-Bermudez *et al*, 2018). Electron acceptors produced by mitochondrial respiration are crucial for aspartate biosynthesis (Sullivan *et al*, 2015), while aspartate treatment reduces ROS production (Quinlan *et al*, 2014). This interplay between aspartate synthesis, the ETC function, and ROS levels indicates a functional loop between the two hypothesized functions of PRDX1.

We next addressed if the aspartate synthesis defect observed in sgPDRX1 cells may impair cell proliferation or survival. Annexin V-Propidium Iodide staining indicated that apoptosis did not increase in U2-OS sgPRDX1 cells when compared to U2-OS WT cells (Fig S5A-B). We then directly tested if sgPRDX1 cells had a growth disadvantage. By mixing U2-OS sgPRDX1 cells with U2-OS WT in equal amounts, we performed a competitive growth assay and determined the percentage of each cell population over a period of 12 days. U2-OS sgPRDX1 cells decreased over time to 30-35% (Fig 5A). However, we observed that the U2-OS sgPRDX1 population recovered PRDX1 expression over time after knock-out generation, probably due to PRDX1 knock-out heterozygous and in-frame deletion selection. Similarly, shPRDX1-treated U2-OS partially restored PRDX1 expression after a few weeks in culture (Fig S5A). Therefore, we repeated the competitive growth assay with a stable HT1080 PRDX1 knock-out clone (PRDX1-/-) (Fig S5A). Here, the percentage of PRDX1-/- cells dropped to 15% (Fig 5B), indicating that the milder effect observed in U2-OS cells was most probably due to population heterogeneity.

**Figure 5:**
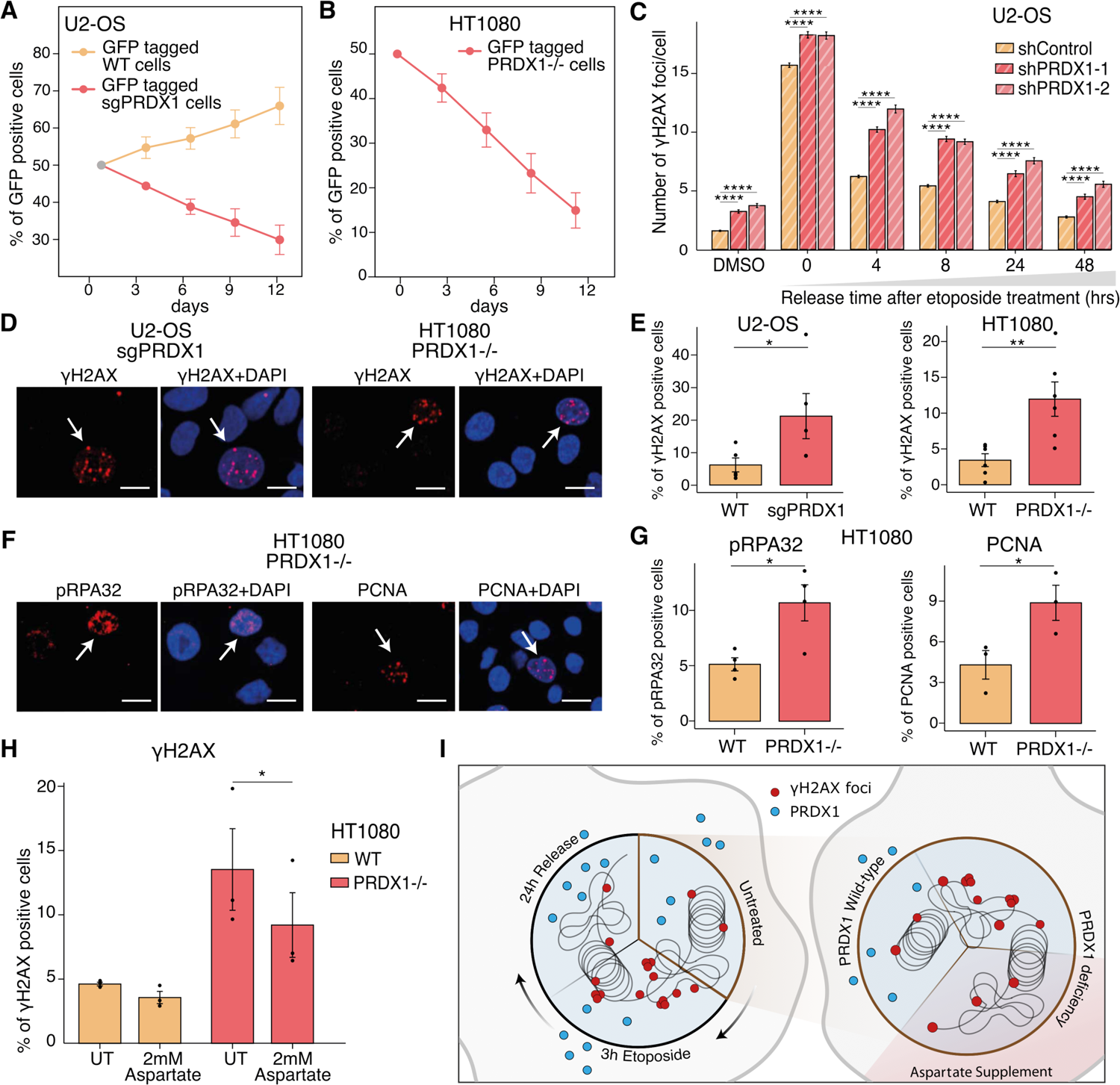
PRDX1 loss generates replication stress that can be partially rescued by exogenous aspartate supplementation. **(A)** Competitive growth assay of U2-OS WT and sgPRDX1 cells. Samples are normalized to day 0, data represent mean and SEM of 3 independent replicates. **(B)** Competitive growth assay of HT1080 WT and PRDX1-/- cells. Samples are normalized to day 0, data represent mean and SEM of 6 independent replicates. **(C)** Kinetics of recovery after etoposide treatment in U2-OS shControl and shPRDX1 cells. Quantification of γΗ2ΑΧ immunofluorescence images represented as mean number of γΗ2ΑΧ foci per nucleus. A minimum of 1,700 cells were quantified for each condition, using CellProfiler, from images acquired with an Opera High Content Screening System. Error bars represent SEM. P-values were calculated using the non-parametric Wilcoxon test. **(D)** Visualization of γΗ2ΑΧ (in red) within DAPI stained nuclei (in blue) in U2-OS sgPRDX1 and HT1080 PRDX1-/- cells. Cells positive for γΗ2ΑΧ are indicated with a white arrow. Cells were untreated and images were acquired on an Opera High Content Screening System, scale bar is 20μm. **(E)** Quantification of images shown in (D). A minimum of 445 cells were quantified for each condition and replicate, using CellProfiler. Data represent mean and SEM of 5 or 6 independent replicates for U2-OS or HT1080 cells, respectively. P-values were calculated using paired t-test. **(F)** Visualization of the replication stress markers pRPA32 and PCNA in HT1080 PRDX1-/- cells. Cells positive for pRPA32 or PCNA are indicated with a white arrow. Cells were untreated and images were acquired on an Opera High Content Screening System, scale bar is 20μm. **(G)** Quantification of images shown in (F). A minimum of 500 cells for pRPA32 staining and 900 cells for PCNA staining were quantified for each condition and replicate, using CellProfiler. Data represent mean and SEM of 4 or 3 independent replicates for pRPA32 or PCNA stainings, respectively. P-values were calculated using paired t-test. **(H)** Quantification of γΗ2ΑΧ foci in HT1080 WT or PRDX1-/- cells, either untreated (UT) or treated for 3 days with 2mM aspartate. A minimum of 2500 cells were quantified for each condition and replicate, using CellProfiler. Data represent mean and SEM of 3 independent replicates. P-values were calculated using paired t-test. **(I)** Scheme of identified roles of PRDX1 in the DNA damage response.

To investigate the underlying mechanisms leading to reduced proliferation, we measured the number of γΗ2ΑΧ foci by high-throughput microscopy in U2-OS PRDX1-depleted cells, in basal conditions, just after induction of DSBs with etoposide, and following recovery in drug-free media (Fig 5C). Cells depleted for PRDX1 had more γΗ2ΑΧ foci in basal conditions and in all other timepoints. No synergy was observed with etoposide treatment and we therefore studied further the role of PRDX1 in the DDR only in basal conditions. It has been recently shown that PRDX1 loss leads to the inhibition of telomerase activity because of increased ROS-induced damage at telomeres (Ahmed & Lingner, 2018). Therefore, we investigated if DNA damage in PRDX1-deficient cells in basal conditions was specifically localized at telomeres. Co-staining of γΗ2ΑΧ with the telomere markers TRF1 and TRF2 in U2-OS sgPRDX1 and HT1080 PRDX1-/- cells, respectively, showed that DNA damage was not restricted to telomeric regions (Fig S5C). We observed that some U2-OS and HT1080 PRDX1-deficient cells significantly accumulated a higher number of γΗ2ΑΧ foci, rather than all cells having slightly more DNA damage than WT cells (Fig 5D-E and S5D). Given that PRDX1 loss impacted aspartate availability, we reasoned that the proliferation defect observed in PRDX1-deficient cells and the accumulation of DNA damage in some cells could be due to reduced nucleotide synthesis that would lead to replication stress. The cellular response to replication stress consists of a cascade of events aiming to limit genome instability, involving numerous proteins. The replication protein A (RPA) is a key protein in this process. This heterotrimer, composed of the subunits RPA70, RPA32 and RPA14, binds and stabilizes single-strand DNA, and is phosphorylated by kinases such as ataxia telangiectasia and Rad3-related protein (ATR) to prevent resection of single-strand DNA (Soniat *et al*, 2019). RPA foci, pRPA and pATR signals are thus markers of replication stress. Additionally, PCNA, an essential replication fork component, regulates the restart of stalled replication forks and therefore also forms foci in replication stress conditions (Essers *et al*, 2005). When analyzing the accumulation of these replication stress markers in HT1080 PRDX1-/- cells, we observed that, similar to γΗ2ΑΧ staining, a subset of cells accumulated high levels of replication stress (Fig 5F-G and S5E). PRDX1-/- cells were twice as likely to beh pRPA or PCNA positive than WT cells (Fig 5F-G). Therefore, the impaired cellular proliferation of PRDX1-deficient cells might in part be explained by replication stress and accumulation of DNA damage.

Finally, since our metabolomics results indicated that PRDX1 loss led to a reduction of aspartic acid and other aspartate-dependent metabolite levels (Fig 4E-F), we questioned whether aspartate supplementation could rescue the observed phenotypes of PRDX1 loss. We quantified the number of γΗ2ΑΧ foci by immunofluorescence after treatment of HT1080 PRDX1-deficient cells with aspartate for 3 days. We observed a reduction in the number of γΗ2ΑΧ positive cells (Fig 5H and S5F). Similarly, the number of pRPA positive cells was reduced after aspartate supplementation, although not significantly, suggesting a decrease in replicative stress (Fig S5G). However, exogenous aspartate supplementation did not rescue the cellular proliferation defect, as measured by competitive growth assay (Fig S5H). This data suggests that the reduced aspartate levels and aspartate-dependent metabolites observed in PRDX1-deficient cells are not the only causes of a proliferative disadvantage. The role of PRDX1 in ROS scavenging, especially at the nuclear level (Fig S3H), might be the principal cause of reduced cellular growth.

Overall, this identifies PRDX1 as an important DNA damage surveillance factor, which is crucial for cellular proliferation. We report that PRDX1 contributes to the clearance of ROS generated in the nucleus following etoposide treatment. In addition, we show that PRDX1 functions in nucleotide homeostasis by maintaining proper intracellular levels of aspartate. Therefore, PRDX1 loss leads to replication stress and accumulation of DNA damage (Fig 5I).

## Discussion

In this study, we took a variety of -omics approaches to evaluate the crosstalk between metabolism and the DNA damage response. By integrating metabolism-focused CRISPR-Cas9 genetic screens, chromatin proteomics and targeted metabolomics in basal conditions and after generation of DSBs by etoposide, we identified metabolic pathways that play a crucial role in maintaining genome integrity. Firstly, several proteins from the ETC were synthetic viable with etoposide treatment and were found to be recruited on chromatin after DSB induction. Nuclear ROS were generated during etoposide treatment and persisted up to 24 hours after treatment. This suggests an important role of the ROS signaling and scavenging processes in maintaining genomic integrity following generation of DNA DSBs with etoposide. Secondly, etoposide treatment induced profound perturbations in the cellular metabolome, which remained altered up to 24 hours after drug release. The main perturbed metabolites were nucleoside-related compounds. This is in line with the literature, since DNA replication is altered following DNA damage, thus modifying nucleotide pools, since nucleotides are necessary to repair DNA damage.

The robustness of our data is confirmed by its intersection with the published literature and the identification of well-known DDR factors in our genetic and proteomic datasets. Data analysis has revealed that many metabolic enzymes and pathways are involved in the generation or the repair of DNA damage, and further investigation is needed to understand how each of them is specifically implicated in the convoluted cellular response to DNA damage, thus opening the door to new research projects. To further dissect which metabolic pathways are involved in DNA repair, the presented datasets could be completed with additional approaches. First, the chromatome proteomics only allows the identification of proteins that directly bind DNA upon DNA damage, while secondary interactors could play a crucial role in the DDR. Therefore, performing mass-spectrometry on nuclear extracts including the soluble nuclear fraction could be a complementary approach and would allow for the discovery of additional metabolic enzymes involved in DNA repair. Similarly, while the panel of metabolites measured in our targeted metabolomics approach is broad and comprises different kinds of metabolites, performing untargeted metabolomics, although more challenging, would allow for the identification of other metabolites perturbed by etoposide treatment, and would facilitate the identification of whole metabolic reactions (Schrimpe-Rutledge *et al*, 2016).

Amongst the metabolic pathways that we identified as linked with the DDR, mitochondrial respiration plays an important role, that could be harnessed to design better anticancer regimens. Indeed, in the genetic screen, genes of the TCA and ETC that are essential for cellular energy production, were synthetic viable with DNA damage, while the HIF complex, which can induce downregulation of mitochondrial respiration, was synthetic lethal. Additionally, in the chromatome dataset, ETC proteins were enriched on chromatin after etoposide treatment, which suggests that these factors participate in the induction of nuclear ROS previously observed following etoposide treatment. Rapidly proliferating cancer cells have an increased demand in biomass synthesis to support cell growth and often face hypoxia due to the lack of oxygenation in tumors (Paredes *et al*, 2021). Therefore, tumors need to undergo metabolic adaptation and change their nutrient utilization during the different stages of malignancy, which can deregulate TCA and ETC processes. Our data suggests that cells with lower ETC activity and heightened glycolytic signaling would be more resistant to induction of DSBs by etoposide treatment. This hypothesis is supported by a new study showing that the cellular metabolism of colorectal cancer cells is activated following treatment with replication stress-inducing drugs, to provide biomolecules necessary for DNA repair and survival (Marx *et al*, 2022). They also discovered that p53-proficient cells upregulate their metabolism more than their p53-deficient counterparts, and therefore rely more heavily on glucose for their survival. Analyzing the metabolic status of the tumors could thus be important to predict patient responses to DNA-damaging agents and design the most appropriate anticancer therapies.

Similarly, cancer cells usually present higher levels of ROS in basal conditions due to their hypermetabolism, but they adapt their antioxidant capacities to maintain redox homeostasis (Kim *et al*, 2019). Recently, anticancer therapies that manipulate ROS levels are being developed, either by inducing more ROS or by inhibiting antioxidant processes, in order to overwhelm cancer cells and disrupt the redox balance, leading to cell death. Prooxidants and antioxidant inhibitors are currently studied in clinical trials, as well as ROS-based repurposed drugs (Wang *et al*, 2021). Our study demonstrates that dual treatment with etoposide and drugs with a similar mode of action could be a potent strategy to kill cancer cells faster and overcome chemoresistance.

Another metabolic pathway that we found tightly connected with DNA damage and repair is nucleotide metabolism. Indeed, deletion of genes involved in this essential cellular process led to accumulation of DNA damage in our genetic screens, probably because of nucleotide pool imbalance and replication stress. In addition, nucleoside-containing metabolites were also drastically perturbed after etoposide treatment in our metabolomics dataset, suggesting that nucleotides were acutely used following DSB generation, which then triggered *de novo* nucleotide synthesis to replenish nucleotide pools. Therefore, we hypothesize that combining etoposide with inhibitors of nucleotide synthesis processes could potentiate the effect of etoposide, by preventing repair of DNA damage. However, nucleotide synthesis is a key cellular process, and the development of inhibitors is limited by side-effect toxicity, which could be alleviated by identifying and targeting regulatory mechanisms specific to tumor cells or to tissue types. Few organ-specific nucleotide metabolases in tumors have been discovered, and development of compounds targeting these enzymes hold great promise for patient treatment (Feng *et al*, 2020; Ma *et al*, 2021).

Intersection of our datasets led to the identification of the peroxiredoxin PRDX1 as a key factor of the DNA damage surveillance processes. This enzyme has a dual function, as a peroxidase with a ROS scavenging function, and as molecular chaperone that can modulate transcription factor activities upon oxidation (Mu *et al*., 2002). It has been shown to have a controversial role in cancer metabolism. On one hand it is overexpressed in some malignant tumors, but on the other hand, PRDX1-deficient mice are prone to develop cancers (Neumann *et al*, 2003). PRDX1 regulates several transcription factors involved in tumorigenesis. For example, it interacts c-Myc oncogene product and suppresses regulation of some target genes, thus limiting tumor growth (Mu *et al*., 2002), but it also regulates NF-κB activity with cytoplasmic PRDX1 suppressing NF-κB activation by preventing peroxide accumulation on one hand, and nuclear PRDX1 enhancing NF-κB activity on the other hand (Hansen *et al*, 2007). The relationship between PRDX1 and cancer depends on many unknown factors and on tissue specificity. Understanding better the functions of this protein is thus of crucial importance.

In our study, we identify two main roles of PRDX1 in the DNA damage response. First, it scavenges nuclear ROS generated by etoposide treatment after translocating to the nucleus. Secondly, it regulates aspartate metabolism, and PRDX1 deletion induces perturbations in aspartate-related metabolites and nucleotide pools. PRDX1 loss severely impacts cellular proliferation and leads to DNA damage and replication stress in basal conditions, which could be due to both accumulation of ROS and alteration of the nucleotide pool. Moreover, it has been shown that targeting PRDX1 sensitizes breast cancer cells to prooxidant agents (Bajor *et al*, 2018). This study shows for the first time that PRDX1 regulates the cellular metabolome and not only the ROS levels. PRDX1 loss affects aspartate metabolism which has consequences on many other metabolites, as aspartate is used for the synthesis of the amino acids arginine and citrulline and plays a crucial role in the *de novo* nucleotide synthesis. Interestingly, while aspartate levels were reduced in PRDX1-deficient cells, several nucleotide monophosphate levels were upregulated. We can hypothesize either that aspartate is over-used to produce nucleotide monophosphates, or that nucleotide salvage pathways are compensating for the downregulated *de novo* nucleotide synthesis due to reduced aspartate. Untargeted metabolomics approaches as well as tracing experiments could help answer this question. Moreover, while nucleotide monophosphates are elevated in PRDX1-deficient cells, nucleotide triphosphates do not follow this trend, perhaps suggesting a defect in the enzymes responsible for this conversion. It has been demonstrated that aspartate metabolism is perturbed in cancer cells to support proliferation. For example, arginosuccinate synthase (ASS1), which converts nitrogen from ammonia and aspartate to urea, is silenced in several cancers, thus leading to an accumulation of cytosolic aspartate and fostering *de novo* pyrimidine synthesis to support cancerous proliferation (Rabinovich *et al*, 2015). The ETC plays an essential role in aspartate synthesis (Birsoy *et al*., 2015). When the ETC is inhibited, for example in hypoxia, a common condition in tumors, aspartate synthesis becomes limiting and cancer cells need to import extracellular aspartate to maintain cellular growth (Garcia-Bermudez *et al*., 2018). In addition, endogenous aspartate is produced in the mitochondria, but needs to be exported in the cytoplasm, where it can be used for nucleotide and amino acid synthesis. In low-glutamine conditions especially, sustaining cytosolic aspartate concentration is critical for cell survival (Alkan *et al*, 2018). Therefore, the controversial role of PRDX1 in cancer metabolism might also be linked with its regulation of aspartate metabolism. In our study, aspartate supplementation of PRDX1-deficient cells did not lead to a full rescue of the phenotypes associated with PRDX1 loss. We observed a decrease in the generation of DNA DSBs in basal conditions after treating HT1080 PRDX1-/- cells with aspartate for 3 days, but we did not observe a rescue on the growth defect. Uptake capacities of exogenous aspartate is cell-type dependent because it requires the presence of specific transporters for cellular import, such as SLC1A3 (Garcia-Bermudez *et al*., 2018), but in most cells endogenous aspartate is preferentially used (Sullivan *et al*, 2018). Analysis of publicly available transcriptomics data (Ghandi *et al*, 2019) indicated that the HT1080 cells used for this experiment only mildly express the transporter SLC1A3 as compared to cell lines with detectable aspartate import activity described by Garcia-Bermudez *et al*. Therefore, exogenous aspartate supplementation might not be sufficient to restore normal levels of aspartate in HT1080 PRDX1-deficient cells. Additionally, the impact of PRDX1 loss on ROS scavenging might be the principal cause of cellular proliferation disadvantage.

Our study sheds light on the interplay between cellular metabolism and the DNA damage response. This is particularly relevant in cancer, which can be considered both a metabolic and a genetic disease and understanding better this crosstalk could help design more efficient and targeted therapies. The implication of PRDX1 in DNA damage and repair processes and in tumorigenesis is clear, but future work will be needed to elucidate in which conditions it functions as a tumor suppressor or on the contrary facilitates tumor development.

## Material and Methods

### 1 Plasmids and reagents

The human metabolic knockout pooled CRISPR library was a gift from David Sabatini (Addgene # 110066). For lentivirus production, the psPAX2 (a gift from Didier Trono; Addgene plasmid # 12260) and VSV.G (a gift from Tannishtha Reya; Addgene plasmid # 14888) packaging plasmids were used. plentiCRISPR v3 was bought from Horizon (Cross *et al*, 2016) and pLKO.2 was a kind gift from Sebastian Nijman (Ludwig Cancer Research, Oxford, UK). For the competition assay, the pKAM-GFP plasmid (a gift from Archibald Perkins, Addgene plasmid #101865) was used to tag the cells with GFP.

Etoposide, NCS from Streptomyces carzinostaticus ≥90% and aspartate were obtained from Sigma-Aldrich.

### 2 Human cell culture

All cells were grown at 37°C at 5% CO_2_ and 3% O_2_. Human bone osteosarcoma epithelial U2-OS cells were purchased from the ATCC cell repository. Human fibrosarcoma epithelial HT1080 cells, both WT and clonal deficient for PRDX1 (PRDX1-/-), were a kind gift from Joachim Lingner (Swiss Institute for Experimental Cancer Research (ISREC), Ecole Polytechnique Fédérale de Lausanne (EPFL)) (Aeby *et al*, 2016). Ablation of protein expression was confirmed by immunoblotting for PRDX1. All cells were cultured in Dulbecco’s Modified Eagle Medium (DMEM, Gibco), supplemented with 10% Foetal Bovine Serum (FBS, Gibco) and 1% penicillin/streptomycin (Sigma-Aldrich).

### 3 Generation of U2-OS PRDX1-depleted cells

For shRNA mediated depletion of PRDX1, two shRNAs (shPRDX1-1 and shPRDX1-2) targeting the coding region of the gene (5’-GATGAGACTTTGAGACTAGTT-3’ and 5’-CCAGATGGTCAGTTTAAAGAT-3’) and one non-targeting shRNA (shControl, 5’-CTTACGCTAGTACTTCGA-3’) were used. The shRNA sequences were obtained from the TRCN database (http://www.broadinstitute.org/rnai/public/gene/search) and cloned into the lentiviral vector pLKO.2 using AgeI and EcoRI restriction sites. For sgRNA mediated depletion of PRDX1, an sgRNA targeting PRDX1 (5’-GCCACAGCTGTTATGCCAGA-3’) was cloned into the lentiviral vector plentiCRISPRv3 using BsmB1 restriction sites. Insertion of shRNA and sgRNA sequences was verified by Sanger sequencing.

Lentiviral particles were produced by transfection of the shRNA-containing pLKO.2 or the sgRNA-containing plentiCRISPRv3 constructs along with the packaging plasmids psPAX2 and VSV.G into HEK-xLenti^TM^ cells (Oxgene) cells using Lipofectamine 2000. Two and three days after transfection, virus-containing supernatant was harvested and centrifuged to remove packaging cells from the supernatant. U2-OS cells were infected by spinfection with the virus-containing supernatant in the presence of polybrene (final concentration 8 ug/mL). Infected cells were selected using puromycin (1.5ug/mL; Sigma-Aldrich) for 72 hr.

To increase knock-out efficiency, sgPRDX1 U2-OS population was additionally nucleofected with the purified S.p. Cas9 nuclease V3 (#1081059, Integrated DNA Technology) together with *in vitro* transcribed sgRNA targeting PRDX1 (5’-GCCACAGCTGTTATGCCAGA-3’). T7 *in vitro* transcription was performed using HiScribe (NEB E2050S), using PCR-generated DNA as template. The 4D-Nucleofector System X-Unit (Lonza) was used for nucleofection, with the SE Cell-Line Solution (V4XC-1032, Lonza) and the CM-104 program, in Nucleocuvette^TM^ strips (Lonza).

Decrease in protein expression in the whole population was confirmed by immunoblotting for PRDX1.

### 4 CRISPR screens

#### a. Pooled CRISPR screen

##### Library amplification

The metabolic CRISPR pooled library was amplified following the distributor’s instructions (Addgene), with a coverage of around 200X.

##### Virus production

HEK-xLenti^TM^ cells (Oxgene) were seeded in 12xT225 flasks 10cm dishes and transfected 24 hours later, with the metabolic CRISPR pooled library, pVSVG and psPAX2 packaging plasmids, using polyethylenimine (PEI) in OptiMeM (Gibco). Medium was changed 10 hours later. 24 and 48 hours later, supernatant containing virus was harvested and centrifuged at 2000 rpm for 5 minutes to remove cell debris. The two batches were pooled together and the virus was concentrated 20X using PEG-8000 and stored at −80°C.

##### Cell infection and harvest

U2-OS cells were spinfected for 3 hours at 2000 rpm and 37°C in 12-well plates with the lentiviral metabolic library at a multiplicity of infection (MOI) of 0.3-0.5 in presence of polybrene (final concentration 8 ug/mL). Immediately after spinfection, cells were collected and seeded in 245mm square dishes with fresh medium. Puromycin-containing medium (1.5 ug/ml) was added the next day to select for transductants. At 7 days post-transduction, cells were re-seeded, and at 9 days post-transduction, they were either treated with 1μM etoposide for 3 hours or left untreated. Treated cells were washed with PBS and released in drug-free media, and after 24 hours release, both untreated and treated cells were harvested. Part of the harvested cells were re-seeded to be harvested at a later timepoint, maintaining 1000X coverage, while the rest of the cells were fixed with ice cold 90% methanol in PBS at a density of 8 million cells/mL, and stored in methanol at −20°C. At 14 days after transduction, both untreated and treated cells were harvested, methanol-fixed and stored at − 20°C.

##### Immunofluorescence staining and FACS

300 million cells harvested at 24 hours of release after etoposide treatment, fixed in methanol and stored at −20°C, were stained for γΗ2ΑΧ and with PI as described in the flow cytometry section, except that γΗ2ΑΧ antibody was diluted 1:300, using 100uL/10 million cells, and AF488-anti mouse antibody was diluted 1:250, using 100uL/10 million cells. After PI staining in batches, cells were filtered and sorted on a SONY SH800 sorter, for the top 5% and the lowest 10% γΗ2ΑΧ populations. Sorted cells of different batches were pooled and stored as cell pellets at −80 until DNA extraction. For consistency, unsorted samples stored in methanol were also washed with PBS, FACS buffer and PBS, and stored as pellets at −80 until DNA extraction.

##### Genomic DNA extraction and sgRNA amplification

Genomic DNA from all samples was extracted using QIAmp DNA Blood Midi kit using a protocol from the Broad Institute, treated with RNaseA and then ethanol precipitated to concentrate the DNA. The sgRNA library was prepared using a one-step PCR with ExTaq polymerase (Takara) and a mixture of P5 forward primers with staggers from 1 to 8 bp and barcoded P7 reverse primers. Cell cycle number was optimized for each sample to ensure that there was no over-amplification and the used DNA input for each sample corresponded to a coverage of ∼500X. PCR products were purified by size-exclusion using magnetic AMPure XP DNA beads (Beckman Coulter) until DNA electrophoresis profiles showed clean peaks (BioAnalyzer 2100, Agilent).

##### NGS analysis

Barcoded samples were pooled in equal quantities after measurement of DNA concentrations by fluorometric quantification (Qubit, ThermoFisher Scientific), and sequenced on one lane of an Illumina HiSeq 3000/4000 machine using single-read sequencing. After de-multiplexing, sgRNA sequences were retrieved by trimming all sequences 5’ to the adapter sequence (5’-GACGAAACACC-3’) and 20 nucleotides 3′ following this. MAGeCK was used for alignment, gRNA count, copy number variation (CNV) correction and gene-level depletion scores (Li *et al*, 2014). MAGeCKFlute (Wang *et al*, 2019) was additionally used to correct for cell cycle-related effects between etoposide-treated and untreated samples. sgRNA counts were normalized to million counts, for each sequencing sample and gene log2(fold-change) was calculated by taking the average of the log2(fold-change) for all sgRNAs targeting the same gene. The next-generation sequencing raw data from this publication have been deposited to the European Nucleotide Archive (ENA) database and assigned the identifier [ERA16463919] (https://www.ebi.ac.uk/ena/browser/view/PRJEB54700).

#### b. Arrayed CRISPR screen

##### Library cloning

The arrayed library was designed to target the top genes whose depletion led to increased γΗ2ΑΧ level at 24 hours release post-etoposide treatment in the pooled metabolic CRISPR screen, excluding the transporters ABCB1 and ABCB7, which have known roles in multidrug resistance. Each gene was targeted by 4 sgRNAs: 2 that were showing the strongest phenotype in the pooled screen and 2 that had the highest score in Toronto KnockOut Library v3 (TKOv3, https://crispr.ccbr.utoronto.ca/crisprdb/public/library/TKOv3/). Additionally, 4 intergenic controls were selected from the pooled library and 3 sgRNAs targeting the DNA repair genes LIG4 and XRCC4 were selected from the TKOv3 library, as positive controls. sgRNAs were cloned in plentiCRISPRv3 using the BsmBI restriction sites, in a 96 well plate format. To amplify the plasmids, Stbl3 bacteria were transformed with the ligation reaction in 96-well deep well plates until OD is approximately 0.1. Then bacteria expressing sgRNAs targeting the same genes were pooled and plasmid DNA extraction was performed using QIAgen Miniprep kit following manufacturer’s instructions. Representation of sgRNAs and cross-contamination between wells was checked by NGS sequencing after one-step PCR to amplify sgRNA sequences, both P5 forward primers and P7 reverse primers being barcoded.

##### Virus production

Virus was produced following the same protocol than for the pooled screen except that HEK-xLenti^TM^ cells (Oxgene) were seeded in 6-well plates and each well was transfected with the mixture of sgRNA-containing plentiCRISPRv3 constructs targeting the same gene using Lipofectamin2000 (ThermoFisher). Virus-containing supernatant was aliquoted and frozen at −80°C.

##### Screen setup

U2-OS cells were spinfected for 2 hours at 2000 rpm and 32°C in 96-well plates with the arrayed library at a high MOI in presence of polybrene (final concentration 8 ug/mL). Puromycin-containing medium (1.5 ug/ml) was added the next day to select for transductants. At 6 days post-transduction, selected cells were seeded in 384 well plates, with duplicated wells for each targeted gene. One day later, they were either treated with 1μM etoposide for 3 hours or 60ng/mL NCS for 1 hour, or left untreated. Treated plates were either fixed with 2% PFA in PBS immediately after treatment or after 20 hours release in drug-free media. Untreated plates were fixed at the same time as 20 hours release plates. γΗ2ΑΧ and DAPI staining was performed as described in the immunofluorescence microscopy section. Images were acquired on an Opera High Content Screening System (Perkin Elmer) using x40 magnification. Quantification of the number of foci per cell was done using CellProfiler software version 4.1.3. To account for inter-experiment variability, the number of foci in each condition was normalized to the number of foci in the untreated intergenic condition for each biological replicate.

### 5 Chromatome proteomics

#### Sample preparation

5 million U2-OS cells were incubated in CHAPS buffer for 20min on ice (0.5% CHAPS in PBS 1x) and centrifuged for 5 min at 720g at 4°C. The supernatant was saved as “Cytoplasmic fraction” and the nuclei resuspended in Cytoplasmic Lysis Buffer (0.1% IGEPAL, 10mM Tris-HCl ph 7, 150mM NaCl). The dirty nuclei were placed on Sucrose Buffer (10mM Tris-HCl ph 7, 150mM NaCl, 25% Sucrose) and centrifuged for 15min at 10000g and 4°C. The nuclei were washed 3 times by resuspending with Nuclei Washing Buffer (0.1% IGEPAL and 1mM EDTA in PBS 1x) and spinning for 5min at 1200g and 4°C. The clean nuclei were resuspended in Nuclei Resuspension Buffer (20mM Tris-HCl ph 8, 75mM NaCl, 1mM EDTA, 50% Sucrose) and lysed by adding Nuclei Lysis Buffer (0.1% IGEPAL, 20mM HEPES pH 7.5, 300mM NaCl, 0.2mM EDTA), vortexing and incubating for 2 minutes on ice. The nuclei extract was centrifuged for 2 min at 16000g and 4°C and the chromatin pellet resuspended in Benzonase Digestion Buffer (0.1% IGEPAL, 15mM HEPES pH 7.5, 5µg/µl TPCK). The chromatin was sonicated on a Bioruptor Pico for 15 cycles 30 sec ON 30 sec OFF in 1.5ml Diagenode tubes, the DNA was digested with 2.5U Benzonase (VWR) for 30 min at RT and the resulting extract saved as “Chromatome fraction”. All buffers contained “Complete” proteinase inhibitor (Roche) according to manufacturer’s directions.

#### Liquid chromatography coupled to tandem mass spectrometry (LC-MS/MS)

The protein concentrations from chromatin enriched samples were determined using the BCA protein assay kit (Applichem CmBH, Darmstadt,Germany), and 10 μg per sample was processed using an adapted Single-Pot solid-phase-enhanced sample preparation (SP3) methodology (Hughes *et al*, 2014). Briefly, equal volumes (125 μl containing 6250 µg) of two different kind of paramagnetic carboxylate modified particles (SpeedBeads 45152105050250 and 65152105050250; GE Healthcare) were mixed, washed three times with 250 µl water and reconstituted to a final concentration of 50 μg/μl with LC-MS grade water (LiChrosolv; MERCK KgaA). Samples were filled up to 100 µL with stock solutions to reach a final concentration of 2% SDS, 100mM HEPES, pH 8.0, and proteins were reduced by incubation with a final concentration of 10 mM DTT for 1 hour at 56°C. After cooling down to room temperature, reduced cysteines were alkylated with iodoacetamide at a final concentration of 55 mM for 30 min in the dark. For tryptic digestion, 400 μg of mixed beads were added to reduced and alkylated samples, vortexed gently and incubated for 5 minutes at room temperature. The formed particles-protein complexes were precipitated by addition of acetonitrile to a final concentration of 70% [V/V], mixed briefly via pipetting before incubating for 18 minutes at room temperature. Particles were then immobilized using a magnetic rack (DynaMag-2 Magnet; Thermo Fisher Scientific) and supernatant was discarded. SDS was removed by washing two times with 200 μl 70% ethanol and one time with 180 μl 100% acetonitrile. After removal of organic solvent, particles were resuspended in 100 μl of 50 mM NH4HCO3 and samples digested by incubating with 2 μg of Trypsin overnight at 37°C. Samples were acidified to a final concentration of 1% Trifluoroacetic acid (Uvasol; MERCK KgaA) prior to immobilizing the beads on the magnetic rack. Peptides were desalted using C18 solid phase extraction spin columns (Pierce Biotechnology, Rockford, IL). Finally, eluates were dried in a vacuum concentrator and reconstituted in 10 µl of 0.1% TFA.

Mass spectrometry was performed on an Orbitrap Fusion Lumos mass spectrometer (ThermoFisher Scientific, San Jose, CA) coupled to an Dionex Ultimate 3000RSLC nano system (ThermoFisher Scientific, San Jose, CA) via nanoflex source interface. Tryptic peptides were loaded onto a trap column (Pepmap 100 5μm, 5 × 0.3 mm, ThermoFisher Scientific, San Jose, CA) at a flow rate of 10 μL/min using 0.1% TFA as loading buffer. After loading, the trap column was switched in-line with a 50 cm, 75 µm inner diameter analytical column (packed in-house with ReproSil-Pur 120 C18-AQ, 3 μm, Dr. Maisch, Ammerbuch-Entringen, Germany). Mobile-phase A consisted of 0.4% formic acid in water and mobile-phase B of 0.4% formic acid in a mix of 90% acetonitrile and 10% water. The flow rate was set to 230 nL/min and a 90 min gradient used (4 to 24% solvent B within 82 min, 24 to 36% solvent B within 8 min and, 36 to 100% solvent B within 1 min, 100% solvent B for 6 min before bringing back solvent B at 4% within 1 min and equilibrating for 18 min). Analysis was performed in a data-independent acquisition (DIA) mode using variable DIA windows. Full MS scans were acquired with a mass range of 375 - 1250 m/z in the orbitrap at a resolution of 120,000 (at 200 m/z). The automatic gain control (AGC) was set to a target of 4 × 10^5^, and a maximum injection time of 54 ms was applied, scanning data in profile mode. A single lock mass at m/z 445.120024 (Olsen *et al*, 2005) was employed. MS1 scans were followed by 41 x MS2 scans with variable isolation windows (variable DIA windows). The MS2 scans were acquired in the orbitrap at a resolution of 30,000 (at 200 m/z), with an AGC set to target 2 × 10^5^, for a maximum injection time of 54 ms. Fragmentation was achieved with higher energy collision induced dissociation (HCD) at a fixed normalized collision energy (NCE) of 35%. Xcalibur version 4.3.73.11 and Tune 3.4.3072.18 were used to operate the instrument. The mass spectrometry proteomics data has been deposited to the ProteomXchange Consortium via the PRIDE partner repository (Perez-Riverol *et al*, 2022) with the dataset identifier [PXD035532] (http://www.ebi.ac.uk/pride/archive/projects/PXD035532). Replicates 4 and 5 were removed from the acquisition as their chromatograms revealed the samples were compromised.

#### Data processing

Chromatin data were batched normalized using ComBat algorithm from the sva R package (version 3.12.0, (Leek *et al*, 2012)) and normalized using the normalize_vsn and median_normalisation functions from the DEP (Zhang *et al*, 2018) and proDA (Ahlmann-Eltze, 2022) packages respectively. The rest of the pipeline was followed according to the DEP package, with the inclusion of impute.mi function for protein-imputation from the imp4p package (Gianetto *et al*, 2020). Known subcellular localization for proteins were obtained from the SubCellularBarCode R package (Arslan, 2021), and the normalization of proteins to their expected whole cell extract (WCE) levels for untreated U2-OS cells was performed through the ProteomicRuler in Perseus and the U2-OS WCE were obtained from the CCLE proteomics dataset (Tyanova *et al*, 2016). Analysis was facilitated by the tidyverse (Wickham *et al*, 2019) collection of packages.

### 6 Metabolomics

#### Sample preparation

U2-OS cells were seeded in 6 well plates. Etoposide treatment (1μM for 3 hours) was performed at different times to be able to terminate the experiment and extract the metabolites simultaneously for all samples. At the last timepoint – treatment for the no release samples – the medium was changed in all wells in order to have growth medium of the same composition at the time of metabolite extraction. Each sample was prepared in triplicates. For metabolite collection, plates containing 0.2-0.4 million cells per well were gently washed with 75mM ammonium carbonate buffer pH 7.4 at room temperature, transferred on ice, and metabolites were extracted with 80:20 ice-cold MeOH:H2O solution. Cells were scraped off and samples were collected in tubes, then snap frozen in liquid nitrogen to stop all metabolic reactions. Once all wells have been collected, samples were thawed and centrifuged in a table-top centrifuge at maximum speed at 4°C. Supernatants containing metabolites were transferred into HPLC vial and stored at −80°C until processing by the metabolomics facility (Pro-Met, CeMM). Cleared extracts were dried under nitrogen. Samples were taken up in MS-grade water and mixed with the heavy isotope labeled internal standard mix.

#### Liquid chromatography coupled to tandem mass spectrometry (LC-MS/MS)

A 1290 Infinity II UHPLC system (Agilent Technologies) coupled with a 6470 triple quadrupole mass spectrometer (Agilent Technologies) was used for the LC-MS/MS analysis. The chromatographic separation for samples was carried out on a ZORBAX RRHD Extend-C18, 2.1 x 150 mm, 1.8 µm analytical column (Agilent Technologies). The column was maintained at a temperature of 40°C and 4 µL of sample was injected per run. The mobile phase A was 3% methanol (v/v), 10 mM tributylamine, 15 mM acetic acid in water and mobile phase B was 10 mM tributylamine, 15 mM acetic acid in methanol. The gradient elution with a flow rate of 0.25 mL/min was performed for a total time of 24 min. Afterwards back-flushing of the column using a 6port/2-position divert valve was carried out for 8 min using acetonitrile, followed by 8 min of column equilibration with 100% mobile phase A. The triple quadrupole mass spectrometer was operated in negative electrospray ionization mode, spray voltage 2 kV, gas temperature 150°C, gas flow 1.3 L/min, nebulizer 45 psi, sheath gas temperature 325°C, sheath gas flow 12 L/min. The metabolites of interest were detected using a dynamic MRM mode.

#### Data processing

The MassHunter 10.0 software (Agilent Technologies) was used for the data processing. Ten-point calibration curves with internal standardization were constructed for absolute quantification of metabolites. Data were analyzed following the DEP R package for differential analysis between conditions and pathway level changes were inferred using the run_mean function from the decoupleR R package (Badia-i-Mompel *et al*, 2022). The metabolomics raw data from this publication have been deposited to the Metabolomics Workbench database (Hughes *et al*., 2014) and assigned the identifier [ST002234] (https://www.metabolomicsworkbench.org/data/DRCCMetadata.php?Mode=Study&StudyID=ST002234).

### 7 Immunoblotting

Cells were lysed in RIPA lysis buffer (New England Biolabs), sonicated and protein concentrations were measured using the Protein Assay Dye Reagent (Biorad). Samples were mixed with NuPAGE LDS Sample Buffer (Invitrogen), boiled for 5 minutes at 98°C and proteins were separated on SDS-PAGE gels and transferred onto Amersham™ Protran nitrocellulose membranes (0.45μm, Cytiva). After 1 hour of blocking in 5% milk in TBS-T (0.1% Tween 20 in 1x Tris-buffered saline), membranes were incubated with primary antibodies at 4°C overnight. Primary antibodies used were against PRDX1 (diluted 1:1,000, ab109506 abcam), Tubulin (diluted 1:10,000, DM1A Cell Signaling), Vinculin (diluted 1:1,000, #13901 Cell Signaling), H3 (diluted 1:10,000, ab1791 Abcam) and FDX1 (diluted 1:500, PA5-59653 Thermo Fisher Scientific). Anti-mouse and anti-rabbit HRP-conjugated goat secondary antibodies (Jackson Immunochemicals) were used at a final dilution of 1:5,000. Immunoblots were imaged using a Curix 60 (AGFA) table-top processor.

### 8 Cellular microscopy

For microscopy-based experiments, U2-OS and HT1080 cells were either seeded in 384-well plates (CellCarrier-Ultra, Perkin Elmer) or on coverslips to assess the subcellular localisation of PRDX1 by confocal microscopy. For staining of pRPA32, RPA32, RPA70, pATR and PCNA, cells were pre-extracted with pre-extraction buffer (10mM PIPES, 100mM NaCl, 3mM MgCl2, 1mM EGTA, 0.5% Triton X-100 and 300mM Sucrose) for 10 minutes at 4°C, followed by Cytoskeleton Stripping Buffer B (10mM Tris pH 7.5, 10mM NaCl, 3mM MgCl2, 1% Tween20, 0.5% sodium deoxycholate) for additional 10 minutes at 4°C (O’Sullivan *et al*, 2021) to only visualise chromatin-bound proteins. All cells were fixed with 2% paraformaldehyde in PBS for 20 minutes at room temperature, washed twice with PBS, permeabilized with 0.5% Triton-X in PBS for 10 minutes at room temperature, washed twice with PBS and blocked for 1 hour with 5% BSA in PBST. Staining with first antibodies (Table S4) was performed overnight at 4°C in 5% BSA in PBST. After three washes with 3% BSA in PBS, staining with mouse-AF568 secondary antibody (diluted 1:2,000, A11004 Molecular Probes) was performed for 1 hour at room temperature. After three washes with 3% BSA in PBS and one wash with PBS, followed by DAPI or 5ug/mL Hoechst 33342 (Life Technologies) staining and washes with PBS, cells were imaged.

Intracellular ROS was measured with CellROX green (Life Technologies), which exhibits bright fluorescence after oxidation and binding to DNA, thus allowing detection of nuclear and mitochondrial ROS, and mitochondria were stained with Mitotracker Deep Red FM (Life Technologies) according to manufacturer’s directions.

Imaging was performed either with an Opera or Operetta High Content Screening System (Perkin Elmer), using the x40 magnification for quantification, an Olympus IXplore SpinSR spinning disk confocal microscope, using the x60 magnification, or a Zeiss LSM700 confocal microscope using the x63 magnification, as indicated in the figure legends. Segmentation of the nuclei using the DAPI or Hoechst channels and quantification of the number of foci per cell or integrated intensity of the nuclear signal was done using CellProfiler software version 4.1.3 or Harmony software, as indicated in the figure legends. Segmentation of the cytoplasm was done based on the Mitotracker signal using the “Find Cytoplasm” option in the Harmony software. When applicable, the threshold to identify positive cells was either the nuclear integrated intensity (pRPA32, pATR) or the number of foci (γΗ2ΑΧ, RPA32, RPA70, PCNA) of the top 5% untreated wildtype cells. Visualization was done with ImageJ Fiji or the Harmony software.

### 9 Flow cytometry

For detection of γΗ2ΑΧ signal, trypsinized cells were fixed in 90% ice cold methanol while vortexing and incubated at least 30 minutes at 4°Cs on a rotation wheel before storage at −20°C or further processing. Cells were then washed with PBS, blocked in FACS buffer (PBS+ 2,5% FBS + 1mM EDTA) and incubated with γΗ2ΑΧ antibody (1:600 in FACS buffer, 100uL/1 million cells) overnight at 4°C on a rotation wheel. After washes with FACS buffer, cells were incubated with AF488-anti mouse antibody (1:600 in FACS buffer, 100uL/1 million cells) for 1 hour at room temperature. After washes with FACS buffer, DNA content was stained by propidium iodide (PI) solution (25ug/mL PI + 200ug/mL RNase A in PBS, 10-20 million cells/mL). Cells were incubated 10-15 minutes at room temperature and stored on ice until flow cytometry acquisition within 3-4 hours.

For determining cell cycle profiles only, methanol-fixed cells were directly washed with PBS and incubated with the PI solution.

For detection of apoptosis, the Pacific Blue™ Annexin V Apoptosis Detection Kit with PI (BioLegend) was used, following manufacturer’s instructions.

Cells were analyzed using a BD LSR-Fortessa X-20. Gating and cell cycle analysis were performed using FlowJo (v10).

### 10 Competitive growth assay

U2-OS WT and sgPRDX1 (population) and HT1080 WT and PRDX1-/- (clone) cells were transduced with pKAM-GFP plasmid and the GFP+ population was sorted using a BD FACSMelody. WT untagged cells and PRDX1-deficient GFP-tagged cells, or the opposite, were mixed together in equal amounts, and percentage of GFP positive cells at day 0 was assessed by analysing an aliquot with flow cytometry. Then cells were harvested and re-seeded every 3 days for 12 days, each time analysing an aliquot with flow cytometry to measure the percentage of GFP positive cells. For the competitive growth assay with aspartate treatment, treated cells were grown in growth medium containing 2mM aspartate from day 0, which was renewed every 3 days. Results were normalized to day 0. Each experiment was performed in technical duplicates or triplicates and biological triplicates.

### 11 Gene ontology-term analysis

Statistical tests for enrichment were performed using the GSEA function in the clusterProfiler R package (Wu *et al*., 2021). To remove redundant terms, due to shared genes, terms were eliminated when they had a high Jaccard Index (larger than 0.3).

### 12 Statistical analysis

Statistical parameters including exact value of n (e.g., total number of experiments, measured cells), deviations, p values and type of statistical test are reported in the respective figure captions. Statistical analysis was performed across biological replicates, by taking the average of the respective technical replicates, when appropriate. Error bars displayed in graphs represent the mean and standard error of the mean (SEM) of at least three biologically independent experiments. Statistical significance was analyzed using paired two-tailed Student’s t-test after testing for normality (Shapiro test) and equal variance (Levene test) or non-parametric Wilcoxon test. p < 0.05 was considered significant. In all cases, ns: not significant (p > 0.05), *: p < 0.05, **: p < 0.01, ***: p < 0.001, ****: p < 0.0001. For the Harmony image quantification, linear regression models were fitted on the log2 integrated intensities to account for both variations in the technical variation between replicates and the biological differences between treatments.

## Data availability

The datasets and computer code produced in this study are available in the following databases:

- CRISPR screen next generation sequencing data: ENA ERA16463919 (https://www.ebi.ac.uk/ena/browser/view/PRJEB54700).
- Chromatome-MS data: PRIDE PXD035532 (http://www.ebi.ac.uk/pride/archive/projects/PXD035532)
- Chromatome-MS analysis code: https://github.com/Skourtis/DIA_Etop_Analysis
- Metabolomics data: Metabolomics Workbench ST002234 (https://www.metabolomicsworkbench.org/data/DRCCMetadata.php?Mode=Study& StudyID=ST002234).

## Author contributions

**Amandine Moretton**: Conceptualization; Investigation; Formal analysis; Methodology; Supervision; Visualization; Writing—original draft; Writing—review and editing. **Savvas Kourtis**: Data curation; Formal analysis; Methodology, Visualization; Writing—review and editing. **Chiara Calabrò**: Investigation; Validation; Writing—review and editing. **Antoni Gañez Zapater**: Investigation; Writing—review and editing. **Frédéric Fontaine**: Investigation; Methodology; Data Curation; Formal Analysis. **André C. Müller**: Methodology; Data Curation; Formal Analysis; Resources. **Sara Sdelci**: Conceptualization; Supervision; Funding acquisition; Writing—original draft; Writing—review and editing. **Joanna I. Loizou**: Conceptualization; Supervision; Funding acquisition; Project administration; Writing—review and editing.

In addition to the <u>CRediT</u> author contributions listed above, the contributions in detail are: AMo, SS and JIL conceptualized the study. SS and JIL obtained funding. AMo, CC and AGZ carried out all cell-based investigations. FF and AMü performed and analyzed the chomatome proteomics experiment. SK performed all bioinformatics investigations. AMo and SK performed analysis and visualization. SS and JIL supervised the study. AMo and SS wrote the original draft and all authors reviewed and edited the final manuscript.

## Acknowledgements

We are thankful to Prof J Lingner (Swiss Institute for Experimental Cancer Research (ISREC), Ecole Polytechnique Fédérale de Lausanne (EPFL)) for providing the HT1080 wild-type and PRDX1-deficient cell lines, and to Prof Sebastian M Nijman (Ludwig Cancer Research, Oxford, UK) for providing the pLKO.2 plasmid. We would like to thank the Biomedical Sequencing Facility (CeMM, Vienna, Austria) for all next generation sequencing and the Metabolomics Facility (CeMM, Vienna, Austria) for the metabolomics analysis. We thank Gerald Timelthaler (Institute of Cancer Research, Imaging Facility) for assistance with the microscopy. We are grateful to Joana Ferreira da Silva and all other members of the Loizou lab, as well as María Lorena Espinar Calvo and all members of the Sdelci lab, for helpful discussions and feedback.

AM and CC were funded by the Austrian Science Fund (grant number P 33024 awarded to JIL). The Loizou lab is funded by an ERC Synergy Grant (DDREAMM Grant agreement ID: 855741). The Sdelci lab’s contributions to this study were funded by an ERC Starting Grant (ERC-StG-852343-EPICAMENTE). This work was funded, in part, by a donation from Benjamin Landesmann. The funder was not involved in the study design, collection, analysis, interpretation of data, the writing of this article or the decision to submit it for publication. CeMM is funded by the Austrian Academy of Sciences.

---

**Supplementary figure 1:**
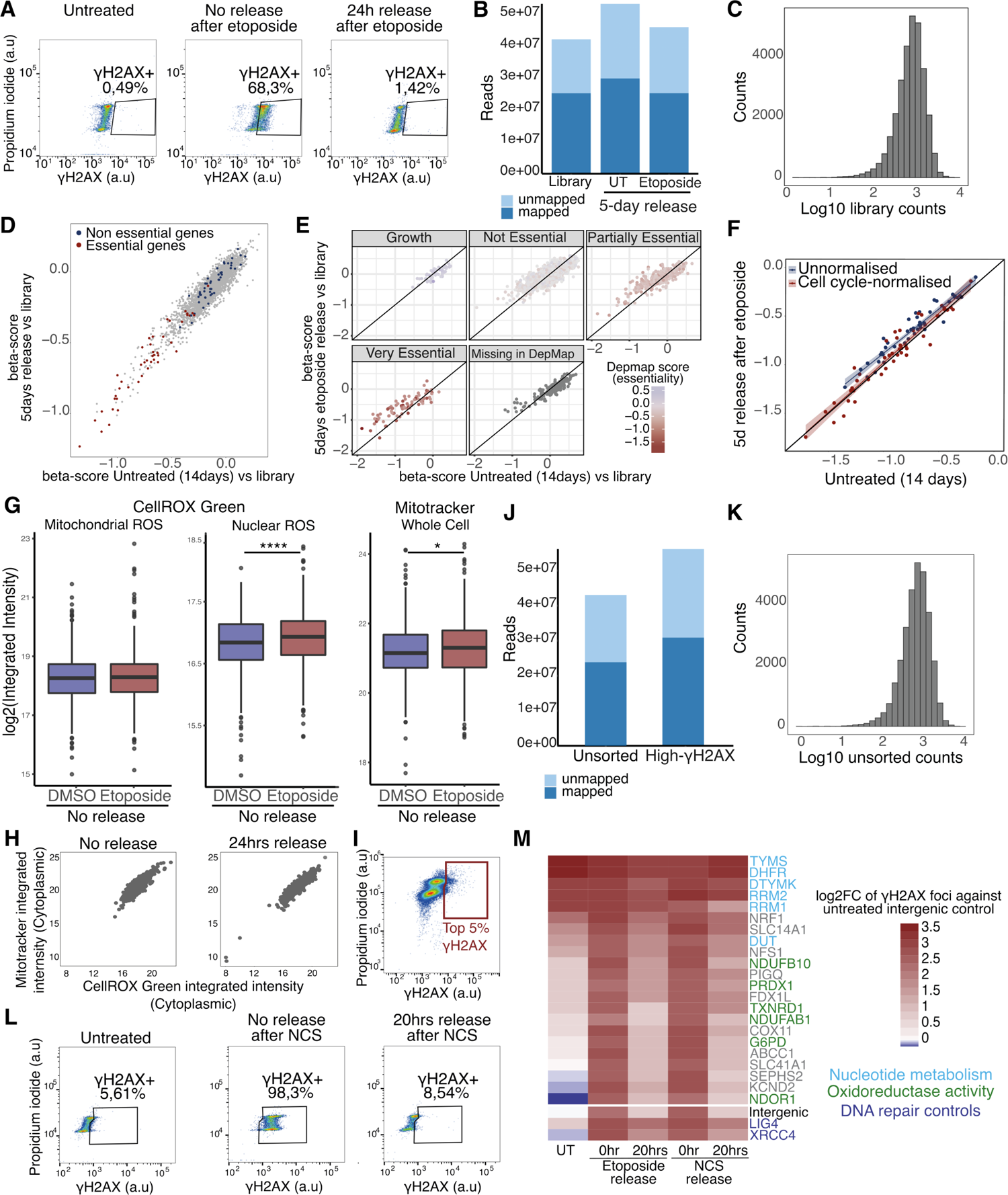
Etoposide-release CRISPR-Cas9 screens with a metabolic library. **(A)** FACS monitoring of γH2AX levels following etoposide treatment and after 24 hours release in drug-free media. **(B)** Mapped reads in the etoposide survival CRISPR screen. **(C)** Distribution of reads in the etoposide survival CRISPR screen. **(D,E)** Comparison of β scores separated by gene essentiality according to MaGECKFlute (D) or DepMap (E). **(F)** MaGECKflute cell cycle normalization based on essential genes. **(G)** Quantification of CellROX Green and Mitotracker stained U2-OS WT cells in DMSO- and etoposide-treated conditions, represented as nuclear or cytoplasmic integrated intensities of CellROX Green signal and whole cell integrated intensities of Mitotracker. A minimum of 1000 cells were quantified for each condition, using Harmony. P-values were calculated using linear regression on the log2 normalized values. **(H)** Correlation of cytoplasmic CellROX Green, representing mitochondrial ROS, and Mitotracker integrated intensities of no release and 24 hours release after etoposide in U2-OS cells. A minimum of 1000 cells were quantified for each condition, using Harmony. **(I)** FACS gating strategy for the high-γH2AX CRISPR screen. **(J)** Mapped reads in the high-γH2AX CRISPR screen. **(K)** Distribution of reads in the high-γH2AX CRISPR screen. **(L)** FACS monitoring of γH2AX levels following NCS treatment and after 20 hours release in compound-free media. **(M)** Quantification of the validation arrayed CRISPR screen using CellProfiler and represented as the mean of the log2 fold change compared to the untreated intergenic control of 3 independent biological replicates.

**Supplementary figure 2:**
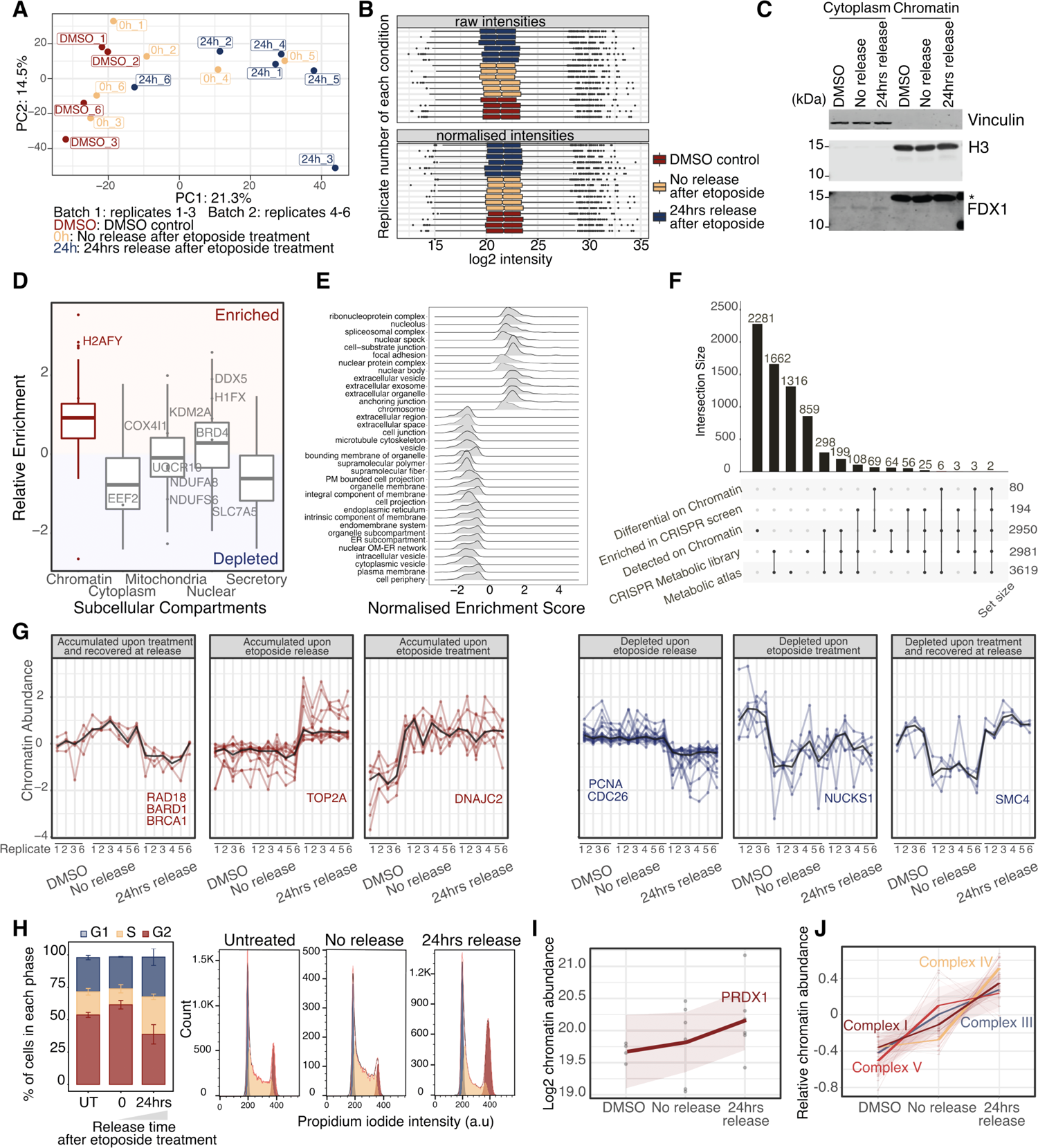
Etoposide-release chromatome proteomics. **(A)** PCA of batch corrected chromatin proteomics samples. **(B)** Median normalization of raw proteomics intensities. **(C)** Western blot confirmation of cell fractionation and chromatin enrichment for one representative replicate (* in FDX1 blot is H3 signal). **(D)** Enrichment of known chromatin proteins normalized to publicly available U2-OS whole cell extract. **(E)** GSEA-Cellular Components of relative enrichment against publicly available U2-OS whole cell extract. **(F)** Overlap of detected and significant genes between the chromatin and CRISPR-Cas9 screen datasets. **(G)** Behavioral clustering of significant proteins with distinct chromatin recruitment patterns. **(H)** FACS monitoring cell cycle profiles after etoposide treatment and release of U2-OS WT cells. **(I)** Kinetics of PRDX1 chromatin recruitment upon etoposide release. **(J)** Relative kinetics of members of ETC complexes chromatin recruitment upon etoposide release. Each protein is centered to its mean value.

**Supplementary figure 3:**
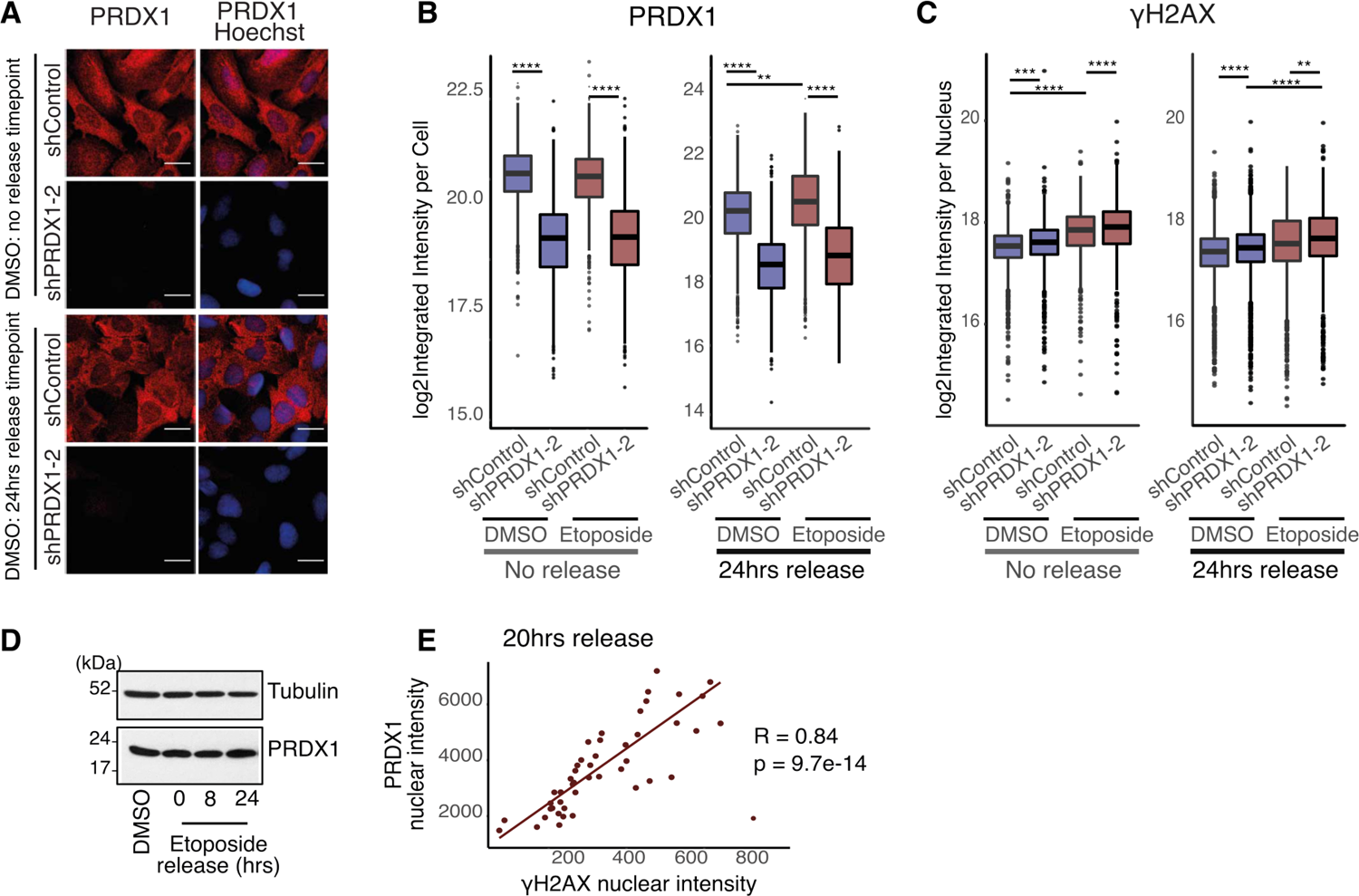
PRDX1 nuclear abundance is related to DNA damage levels. **(A)** Visualization of PRDX1 (in red) and Hoechst (in blue) in U2-OS shControl and shPRDX1 cells in DMSO-treated conditions. Images were acquired with the Operetta High Content Screening System. Scale bar is 25μm. **(B-C)** Quantification of whole cell (B) and nuclear (C) integrated intensity of PRDX1 (B) and γH2AX (C) immediately after etoposide treatment and at 24 hours release compared to DMSO control, in U2-OS shControl and shPRDX1 cells. A minimum of 1000 cells were quantified for each condition, using Harmony. P-values were calculated using linear regression on the log2 normalized values. **(D)** Immunoblot showing PRDX1 abundance in total extracts of U2-OS cells after etoposide treatment and release in drug-free media. Tubulin is used as a loading control. **(E)** Correlation between γΗ2ΑΧ and PRDX1 nuclear integrated intensities in U2-OS WT cells at 20 hours etoposide release.

**Supplementary figure 4:**
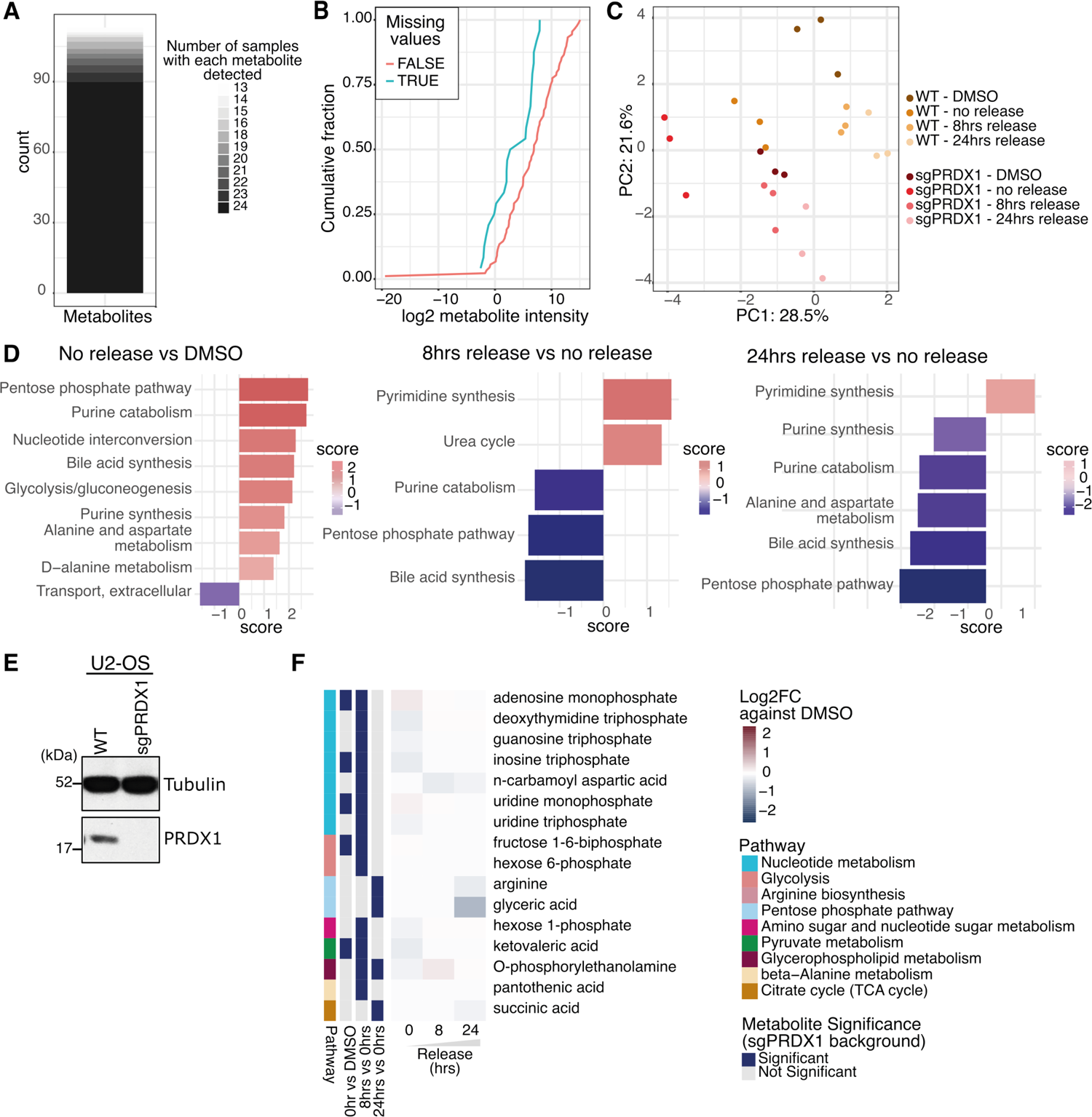
Etoposide-release metabolomics. **(A)** Number of metabolites detected per sample in the targeted metabolomics experiment. **(B)** Distribution of intensities of consistently detected- and partially detected-metabolites showing an intensity-dependent detection pattern. **(C)** PCA plot for all samples in the metabolomics experiment. **(D)** Significantly affected metabolic pathways upon etoposide treatment and release in U2-OS WT cells. **(E)** Immunoblot for PRDX1 and Tubulin on protein extracts from U2-OS WT and sgPRDX1 cells used in the metabolomics experiment. **(F)** Significantly affected metabolites due to etoposide treatment and 24 hours release in U2-OS sgPRDX1 cells.

**Supplementary figure 5:**
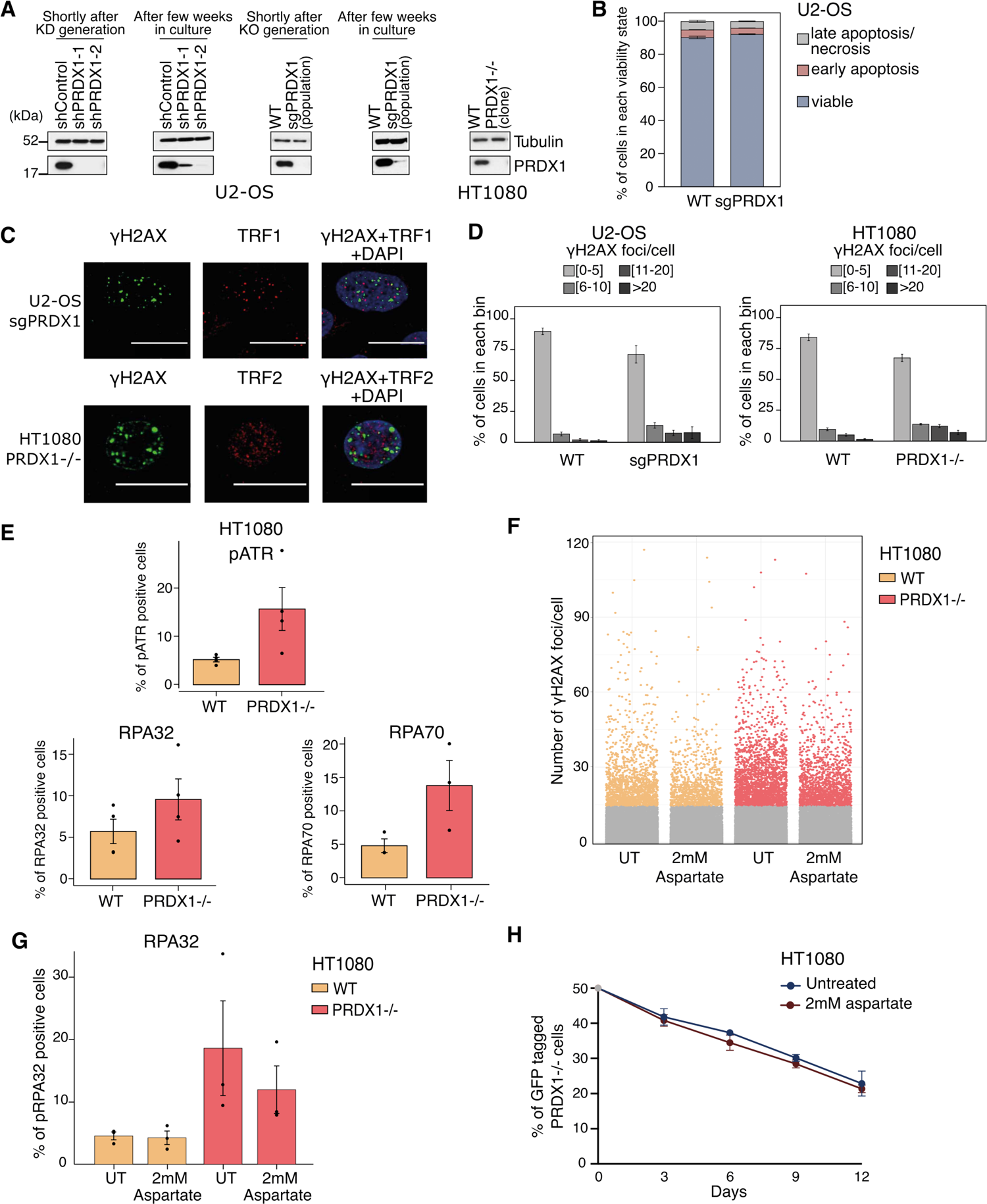
PRDX1 loss generates replication stress that can be partially rescued by exogenous aspartate supplementation. **(A)** Immunoblot for PRDX1 and Tubulin on protein extracts from U2-OS WT, sgPRDX1, shControl and shPRDX1 cell populations, and HT1080 WT and PRDX1-/- cells. **(B)** Detection of apoptosis in U2-OS WT and sgPRDX1 cells using Annexin V-Propidium Iodide staining. Data represent mean and SEM of 3 independent replicates. **(C)** Visualization of the DNA damage marker γΗ2ΑΧ (in red) and the telomere markers TRF1 or TRF2 (in green) in U2-OS sgPRDX1 or HT1080 PRDX1-/- cells, respectively. Cells were untreated and images were acquired on an Olympus spinning disk confocal microscope, scale bar is 20μm. **(D)** Quantification of images shown in Fig 5E, represented as the percentage of cells in each bin of γΗ2ΑΧ foci number. A minimum of 445 cells were quantified for each condition and replicate, using CellProfiler. Data represent mean and SEM of 5 or 6 independent replicates for U2-OS or HT1080 cells, respectively. **(E)** Quantification of immunofluorescence images after staining of the replication stress markers RPA32, RPA70 and pATR, in HT1080 WT and PRDX1-/- cells. A minimum of 375 cells for RPA32 staining, 900 cells for RPA70 staining and 180 cells for pATR staining were quantified for each condition and replicate, using CellProfiler. Data represent mean and SEM of 4 independent replicates for RPA32 and pATR stainings and 3 independent replicates for RPA70 staining. P-values were calculated using paired t-test. **(F)** Quantification of γΗ2ΑΧ foci in HT1080 WT or PRDX1-/- cells, either untreated (UT) or treated for 3 days with the 2mM aspartate. 3 independent replicates are represented. **(G)** Quantification of pRPA32 nuclear integrated intensity in HT1080 WT or PRDX1-/- cells, either untreated (UT) or treated for 3 days with 2mM aspartate. A minimum of 300 cells were quantified for each condition and replicate, using CellProfiler. Data represent mean and SEM of 3 independent replicates. P-values were calculated using paired t-test. **(H)** Competitive growth assay of HT1080 WT and PRDX1-/- cells after 2 mM aspartate treatment. Samples are normalized to day 0, data represent mean and SEM of 3 independent replicates.

## Supplementary tables and data (Annex)

**Table S1** Gene lists belonging in each of the categories of Fig S2F.

**Table S2** Proteins found on chromatin which were significantly enriched/depleted in CRISPR screens.

**Table S3** Genes causing the KEGG-based GSEA enrichment of ‘Chemical Carcinogenesis-ROS’ term, and the ETC complex they belong in.

**Table S4** References and dilutions of antibodies used in immunofluorescence microscopy experiments.

**Source Data_1** Beta scores for etoposide survival and high-γH2AX CRISPR screens.

**Source Data_2** Differential abundant proteins on chromatin of DMSO-treated, etoposide-treated or 24 hours-etoposide release in U2-OS cells.

**Source Data_3** Normalized and imputed metabolite abundances for U2-OS WT and PRDX1-deficient cells, upon etoposide treatment and release.

